# *Plasmodium falciparum* transmission in the highlands of Ethiopia is driven by closely related and clonal parasites

**DOI:** 10.1101/2023.06.09.544365

**Authors:** Aurel Holzschuh, Yalemwork Ewnetu, Lise Carlier, Anita Lerch, Inna Gerlovina, Sarah Cate Baker, Delenasaw Yewhalaw, Werissaw Haileselassie, Nega Berhane, Wossenseged Lemma, Cristian Koepfli

**Affiliations:** Department of Biological Sciences, Eck Institute for Global Health, University of Notre Dame, Indiana, USA; University of Gondar, Gondar, Ethiopia; Trinity College Dublin, Dublin, Ireland; Noul Inc, Seoul, Republic of Korea; EPPIcenter research program, Division of HIV, ID and Global Medicine, Department of Medicine, University of California, San Francisco, CA, USA; Tropical and Infectious Disease Research Center, Jimma University, Jimma, Ethiopia; School of Public Health, Addis Ababa University, Addis Ababa, Ethiopia

**Keywords:** molecular malaria surveillance, *Plasmodium falciparum*, population genetics, molecular diagnostics, genomic surveillance, amplicon sequencing, drug resistance, pfhrp2, pfhrp3, rapid diagnostic test (RDT) molecular epidemiology, next generation sequencing (NGS)

## Abstract

Malaria cases are frequently recorded in the Ethiopian highlands even at altitudes above 2,000 m. The epidemiology of malaria in the Ethiopian highlands, and in particular the role of importation by human migration from the highly endemic lowlands is not well understood. We characterized the parasite population structure and genetic relatedness by sequencing 159 *P. falciparum* samples from Gondar and an additional 28 samples from Ziway using a highly multiplexed droplet digital PCR (ddPCR)-based amplicon deep sequencing method targeting 35 microhaplotypes and drug resistance loci. Diversity was moderate (mean H_E_: 0.54), and infection complexity was low (74.9% single clone infections). A significant percentage of infections shared genomic haplotypes, even across transmission seasons, indicating persistent local and focal transmission. Multiple clusters of clonal or near-clonal infections were identified, highlighting the overall high genetic relatedness. Frequently, infections from travelers were the earliest observed cases, suggesting that parasites may have been imported and then transmitted locally. We observed population structure between Gondar and Ziway, although some haplotypes were shared between sites. 31.1% of infections carried *pfhrp2* deletions and 84.4% *pfhrp3* deletions, and 28.7% *pfhrp2*/*pfhrp3* double deletions. Parasites with *pfhrp2/3* deletions and wild-type parasites were genetically distinct. Mutations associated with resistance to sulfadoxine-pyrimethamine and lumefantrine were observed at near-fixation, but no mutations in *pfk13* were found. In conclusion, genomic data corroborates local transmission and the importance of intensified control in the Ethiopian highlands.

## INTRODUCTION

Despite significant reductions in malaria morbidity and mortality, malaria remains a major public health problem worldwide, causing 247 million cases and killing more than 600,000 people in 2021 (1). In many countries, areas of low malaria transmission that could be targeted for elimination are close to areas of higher transmission. In Ethiopia, approximately 60% of the population is at risk of malaria, with *Plasmodium falciparum* being responsible for the majority of malaria cases and deaths (2,3). Transmission is thought to occur mainly at altitudes below 2,000 m (4,5). The highlands are considered regions of “unstable” malaria transmission (6,7). Migration from the highlands to the lowlands is common, either for short-term visits or to work on large farms (8–10). Highly malaria-endemic lowland areas may serve as a source of malaria transmission to low malaria-endemic highland areas. Travel has been implicated as a major risk factor for *P. falciparum* infection in low-transmission areas in Ethiopia (5,11), and the return of travelers to the highlands has been associated with increased numbers of malaria infections (8–10). However, the potential contribution of returnees to malaria transmission in the highlands is not clear.

Malaria parasite genetic data can be useful for identifying foci of sustained transmission (i.e., sources and sinks of infection) (12–14), estimating connectivity between parasite populations (e.g., between different regions of a country) (15), distinguishing local from imported cases (12), tracking the spread of drug resistance markers (16–19), and assessing transmission intensity (20–22). A large population genomic study found that Ethiopia has a distinctive and highly divergent *P. falciparum* population compared to other regions of Africa (23). Previous studies generally described low proportions of complex infections (i.e., multiplicity of infection, MOI) and moderate population-level genetic diversity in Ethiopia (23–25), suggesting infrequent recombination and smaller parasite population sizes in some areas. Varying prevalence of mutations in the drug-resistance loci *pfdhps*, *pfdhfr, and pfmdr1* have been observed across Ethiopia (25–29), some of which have reached fixation or near-fixation in certain locations.

Most rapid diagnostic tests (RDTs) used to diagnose falciparum malaria detect the histidine-rich proteins 2 and 3 (HRP2 and HPR3), encoded by the *pfhrp2* and *pfhrp3* genes. However, parasites lacking these genes can evade HRP2-based RDT detection (30). A high prevalence of *pfhrp2/3*-deleted parasites has been reported in the Horn of Africa, including Ethiopia (31–38).

Genetic data on *P. falciparum* parasites from Ethiopia remain scarce, and no data are available on parasite diversity, connectivity between highland and lowland populations, drug resistance, and *pfhrp2/3* deletion status from the highland area of Gondar, and Ziway. We used a high-throughput multiplexed amplicon deep sequencing (AmpSeq) method targeting highly diverse microhaplotypes and drug resistance loci previously used in a low-transmission setting in Zanzibar (39). Here, we characterize for the first time the parasite population structure across the high and low transmission seasons, map antimalarial and diagnostic resistance profiles, and define parasite genetic relatedness.

## METHODS

### Study sites

For the current study, samples collected from Gondar in the northwestern highlands were sequenced, along with a smaller set of samples collected from Ziway, located approximately 550 km south of Gondar in central Ethiopia, for comparison.

Gondar is a town of approximately 200,000 people at about 2100 meters above sea level in the Amhara Region, where malaria cases are reported in almost all districts (4,40,41). Of all the Western, Northern and Central zones of Gondar, the lowland districts are highly malaria endemic, while the remaining districts are characterized by low to moderate malaria transmission (4,40,41). There are ongoing reports of cases of both *P. falciparum* as the predominant species and *P. vivax* in this area (42), but data are scarce. Similar to reports from other highland areas in Ethiopia, malaria transmission is highly seasonal with annual fluctuations (42). The main rainy season with the highest number of malaria cases is from October to January, with a shorter rainy season around June-July. Travel to the lowlands is common. Many individuals travel to the lowlands to work on agricultural farms and return to highland areas of Gondar in November and December after the harvest and in time for Orthodox Christmas. The importance of parasite importation by returning travelers on local transmission is unknown.

A total of 2,079 blood samples from all age groups were collected in the Maraki Health Clinic in Gondar, between November 19, 2019, and September 22, 2020 (Figure 1). The Maraki Health Clinic is the largest health center in the area. Additional patient information including age, sex, recent treatment, and recent travel history within the past month was collected. At least two dried blood spot (DBS) samples (50 µL per spot) were collected on Whatman 3MM filter paper (GE Healthcare Life Sciences). DBS samples were placed in plastic bags and stored at 4°C until processing. A total of 206/2,079 (9.9%) samples were *P. falciparum*-positive by microscopy. This number represents the vast majority of all cases during the study period, with only short periods when samples were not collected due to unavailability of study personnel. DBS was available for 198 (96.1%) of the microscopy-positive *P. falciparum* specimens.

**Figure 1.**
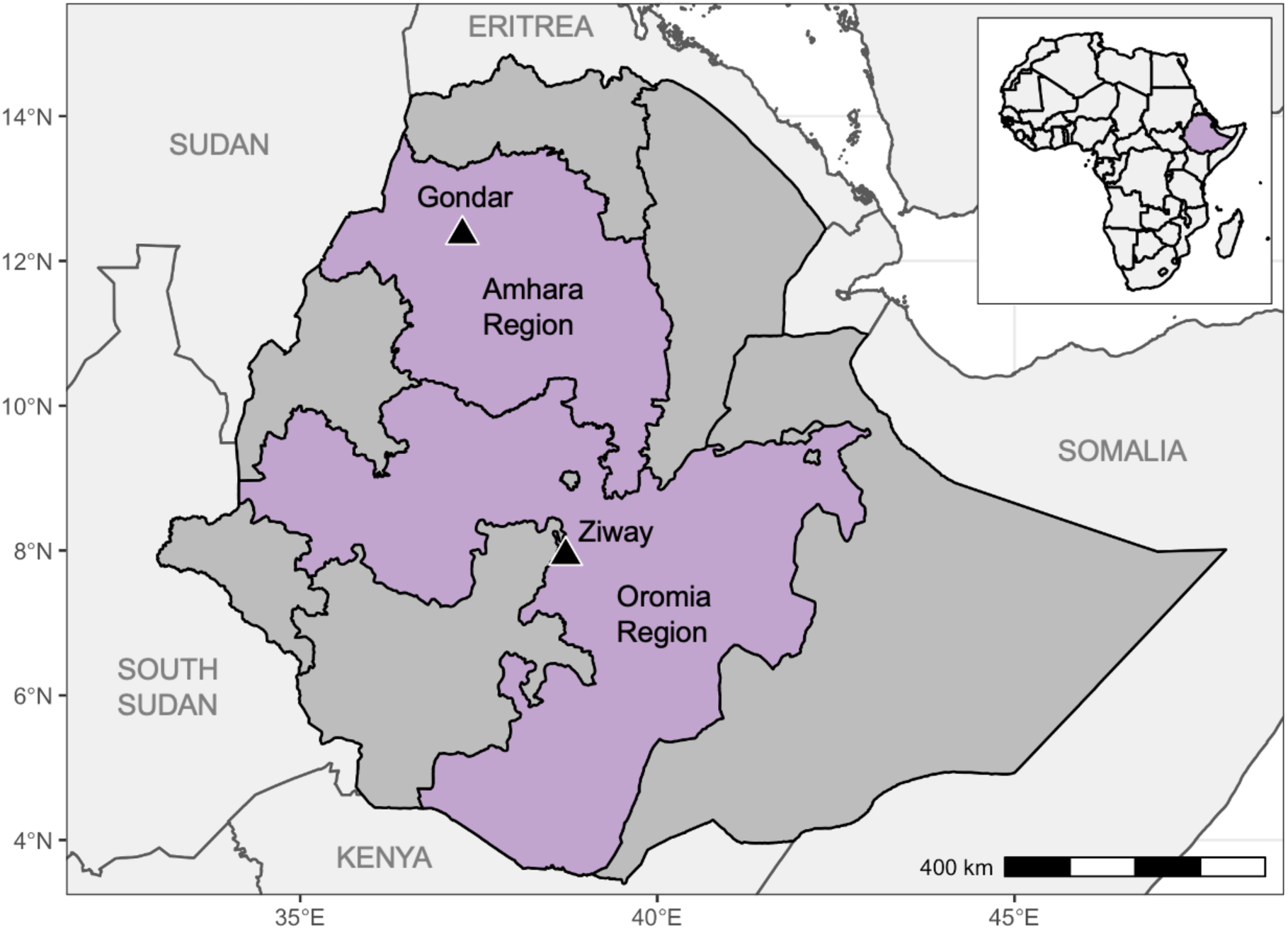
Map of Ethiopia. Triangles show the locations of the Maraki Health Clinic in Gondar and the Batu and Dembel Health Centers in Ziway. Amhara and Oromia regions are highlighted in purple.

Ziway in the Oromia Region is located at an altitude of approximately 1600 m. A total of 1,920 clinical samples were collected from individuals presenting with febrile illness at Batu and Dembel Health Centers in June and July 2019 (43). A total of 56/1,920 (2.9%) samples were *P. falciparum*-positive by microscopy and were included in the current study. Ziway is home to large flower farms that attract workers from across the country. Travel data were not available for samples from Ziway.

In both study sites, malaria is routinely diagnosed by microscopy in health centers and hospitals, but RDTs are used by health extension workers.

### DNA extraction and multiplex amplicon deep sequencing

From 198 *P. falciparum* microscopy-positive patients with a DBS available from Gondar, DNA was extracted from 50 μL DBS and eluted in 50 μL, as previously described (44). DNA from the 56 *P. falciparum* microscopy-positive DBS samples from Ziway was extracted previously (43). For all samples extracted from DBS, parasite density was quantified using *varATS* ultra-sensitive qPCR (45) in a total volume of 12 μL, including 4 μL DNA, corresponding to 4 μL blood. All samples with >1 parasite/µL of blood were genotyped. Of the 198 DBS from Gondar, 182 samples were confirmed *P. falciparum* positive by qPCR, of which 168 were genotyped using a highly multiplexed amplicon deep sequencing protocol targeting 28 microhaplotypes and 7 drug-resistance loci as described previously (39) (Supplementary Tables 1 – 4). Of the 56 *P. falciparum*-positive DBS samples from Ziway, 34 had >1 parasites/µL and were sequenced.

Multiplexed amplification in microdroplets was performed as previously described (39). The final library at a concentration of 1.6pM, including 25% *PhiX* v3 control spiked in, was sequenced on a NextSeq 500 instrument (Illumina) using 150 bp paired-end sequencing with dual indexing using NextSeq at the University of Notre Dame Genomics and Bioinformatics Core Facility. Barcode sequences are provided in Supplementary Table 5 and Additional file 1. Targeted amplicon deep sequencing data was analyzed using the *HaplotypR* version 0.3 bioinformatic pipeline (https://github.com/lerch-a/HaplotypR), using filter criteria defined and explained previously (46). Haplotype calling required a within-host haplotype frequency of ≥1% and a minimum coverage of ≥50 reads per haplotype in ≥2 samples. A total of 187 samples (Gondar *n*=159; Ziway *n*=28) with data in at least 18 loci (i.e., >50% coverage) were included in the final analyses. Haplotype calls and metadata are available in Additional file 2 and 3.

### Multiplicity of infection and population level genetic diversity

Multiplicity of infection (MOI) was defined as the highest number of alleles detected by at least two of the 28 microhaplotype loci. Population level genetic diversity was estimated by expected heterozygosity (H_E_) using the formula, 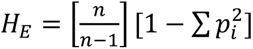, where *n* is the number of samples genotyped and *p_i_* is the frequency of the *i^th^* allele in the population. Drug resistance loci were excluded from MOI and H_E_ calculation. MOI and H_E_ were compared by transmission season and travel history.

### Population genetic analyses

Several methods were used to detect population genetic structure. Principal component analysis (PCA), using the dominant alleles, was performed using the *ade4* (47) package. Discriminant analysis of principal components (DAPC) was done using the *adegenet* (48) package. To determine the appropriate number of principal components to retain for DAPC and to avoid overfitting of the discriminant functions, alpha-score optimization was used. We inferred *P. falciparum* population structure (from dominant alleles) using Bayesian clustering package *rmaverick* v1.1.0 (49), under the non-admixture model with 20,000 burn-in iterations, 10,000 sampling iterations and 50 thermodynamic rungs. As a measure of inbreeding in each population, we computed the multilocus linkage disequilibrium (mLD) using LIAN version 3.7, applying a Monte Carlo test with 100,000 permutations (50). Only samples with complete haplotypes were included.

### Pairwise genetic relatedness by identity by descent (IBD)

We used the R package *dcifer* version 1.2.0 (https://cran.r-project.org/web/packages/dcifer/index.html), an IBD-based method to estimate pairwise genetic relatedness between polyclonal infections (51). *Dcifer* allows for unphased multiallelic data, such as microhaplotypes, and provides likelihood-ratio *p-*values adjusted for one-sided tests. Briefly, naïve MOI estimates and subsequently estimates of population allele frequencies adjusted for MOI were calculated from the samples. We used likelihood-ratio test statistic to test the null hypothesis that two samples are unrelated (*H_0_:* IBD = 0) at significance level *α* = 0.05 (with the procedure adjusted for a one-sided test). We assessed mean IBD and the number of related samples for different groups and performed permutation tests (100,000 permutations) to determine which combinations had higher mean IBD and more related samples than expected by chance. Relatedness networks of parasites were created using the R *igraph* package.

### Pfhrp2/3 deletion typing

Samples that were successfully sequenced (*n*=187) were typed for *pfhrp2* exon 2 and *pfhrp3* deletion by droplet digital PCR (ddPCR) (52). The assay quantifies *pfhrp2* or *pfhrp3* and a control gene (*serine-tRNA ligase*) in a single tube with very high specificity, thus providing highly accurate data on deletion status (52). The assay targets parts of *pfhrp2* exon 2 that encodes for the antigen detected by the RDT. The targeted *pfhrp2* exon 2 region was deleted in all reported cases where *pfhrp2* deletion breakpoints were studied (30,53). The following criteria were used for the analysis of the ddPCR data: A minimum of ten droplets positive for *tRNA* was required to include a sample in the data analysis. Infections were considered “mixed” for either *pfhrp2* or *pfhrp3* gene if the ratio between *pfhrp2/3* and *tRNA* was between 0.1 – 0.6 (i.e., >0.6 was considered *pfhrp2/3+* and <0.1 was considered *pfhrp2/3*-), as previously determined (52). No RDT screening was performed on any of the samples.

To explore genetic differentiation (from dominant alleles) between parasite populations we calculated *G*^′′^ (54), a modified version of Hedrick’s *G^′^* that corrects for a bias stemming from sampling a limited number of populations, for *pfhrp2/3*-deleted parasites versus *pfhrp2/3* wildtype parasites. Briefly, *G*^′′^ was calculated using the formula: 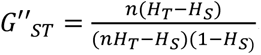, where *H_T_* and *H_S_* are the total and the subpopulation heterozygosity, respectively, and *n* is the number of populations sampled. We used permutation testing to determine the statistical significance of *G*^′′^ values (10,000 permutations).

### Analysis of drug resistance loci

We measured the prevalence of five drug resistance markers: *Pfdhfr* (PF3D7_0417200; amino acids 50, 51, 59, 108 and 164)*, Pfdhps* (PF3D7_0810800; amino acids 540, 581 and 613)*, Pfmdr1* (PF3D7_0523000; amino acids 86 and 184), *Pfmdr2* (PF3D7_1447900; amino acids 484 and 492), and *Pfk13* (PF3D7_1347700; amino acids 481, 493, 527, 537, 538, 539 and 543). The frequency of mutations in the population was calculated as the number of samples containing mutations over the total number of samples genotyped. The *UpSetR* package in R was used to visualize different combinations of multidrug-resistant parasites (monoclonal samples only).

### Public whole genome sequencing data

Fastq files from 198 monoclonal whole genome sequencing (WGS) samples (F_WS_ >0.95) from 16 different countries included in the MalariaGEN *Plasmodium falciparum* Community Project were downloaded from the European Nucleotide Archive using fasterq-dump (v.2.10.8) and sample accession numbers on 23 February 2022 (Supplementary Table 6). Microhaplotype sequences were extracted using BWA-MEM (57), SAMtools (58), and BEDTools (59) software. Briefly, reads were aligned to the *P. falciparum* 3D7 v3 reference genome assembly using ‘bwa mem’. We then used *samtools* library ‘sort’, ‘index’ and ‘view’ functions to extract microhaplotype regions. The *bcftools* library ‘mpileup’, ‘view’ (DP>=20, MQ>=30), ‘filter’ (%QUAL<99) and ‘consensus’ functions, and *bedtools* ‘getfasta’ were used to extract the final microhaplotype sequences (Additional file 4). We inferred genetic differentiation by DAPC and genetic relatedness was estimated using *dcifer* (51).

## RESULTS

### Study population and amplicon deep sequencing results

187/202 (92.6%) sequenced DBS samples had data in ≥18 loci (>50% coverage) and were included in the analyses (159 from Gondar, 28 from Ziway, Supplementary Figures 1a and 2). The 187 samples achieved high coverage of loci (median 100%, IQR 97.1 – 100) (Supplementary Figure 1b), with some variation in the average number of reads per target (Supplementary Figure 1c). 31/35 markers had ≥50 reads in at least 90% of samples (Supplementary Figure 1d). Among successfully sequenced samples from Gondar, 10/159 (6.3%) patients reported recent travel within the past month, 63 were collected in the wet season from November to January, and 96 in the dry season from February to September (Supplementary Figure 2). No travel data were available for samples from Ziway.

### Multiplicity of infection and population diversity

Polyclonal infections were found in only 25.1% of infections, with higher mean MOI in Gondar than in Ziway (Table 1, Figure 2a). In Gondar, we observed some subtle trends in MOI between transmission seasons (Table 1). Population-level diversity was comparable between Gondar and Ziway (mean H_E_: 0.53 vs. 0.50, *p* = 0.359; Table 1, Figure 2b). In Gondar, mean heterozygosity was significantly higher in the dry season compared to the wet season (Table 1). Overall, the microhaplotypes showed moderate genetic diversity (mean H_E_: 0.54, range: 0 - 0.78) in our study sites; 21 of the 28 microhaplotypes had H_E_ ≥0.5 (Supplementary Figure 3). Compared to the expected global heterozygosity (mean H_E_: 0.67, range: 0 - 0.95) of 198 monoclonal samples from 16 different countries, the observed heterozygosity of microhaplotypes in this study was significantly lower (0.54 vs 0.67, *p* = 0.014) with low correlation between the global and local heterozygosity (Pearson’s *r* = 0.21; *p* = 0.196), highlighting that many microhaplotypes are less diverse in Ethiopia than in the global population. (Figure 3c).

**Figure 2.**
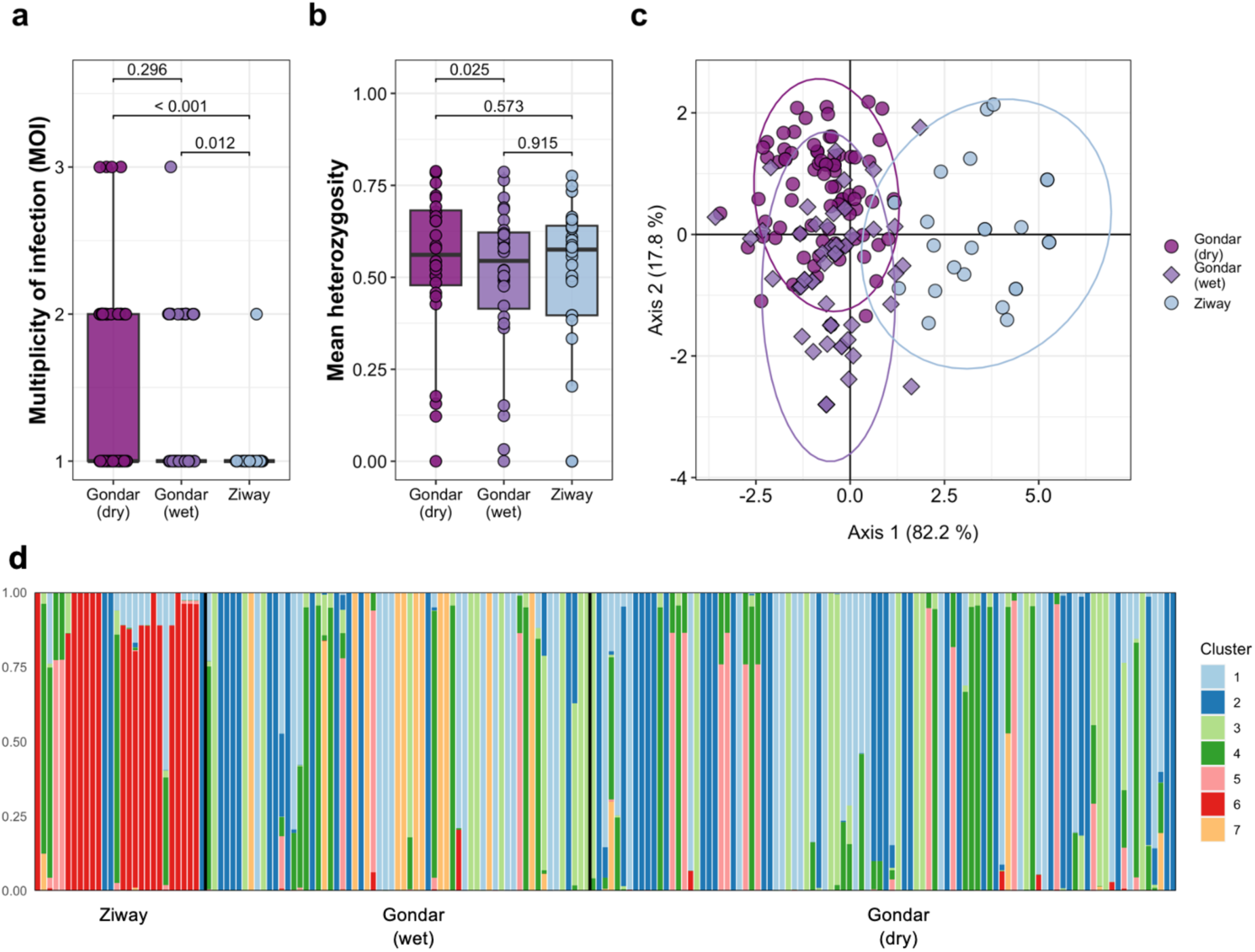
Genetic diversity and population structure in Ethiopia. **A**, Multiplicity of infection (MOI) of samples from Gondar (n=159) separated by dry (n=96) and wet (n=63) season, and Ziway (n=28). **B**, Mean heterozygosity of 28 microhaplotypes in Gondar and Ziway. Bonferroni adjusted pairwise p-values are indicated. **C,** Separation of P. falciparum samples using discriminatory analysis of principal components (DAPC) of the 28 microhaplotypes. The first 31 PCs are shown. Appropriate number of PCs to retain for DAPC (so as to avoid over-fitting) was determined by alpha score optimization. Each dot represents a sample colored by study site (and transmission season for Gondar). **D**, Population cluster analysis of P. falciparum microhaplotypes from dominant alleles in Gondar and Ziway. Individual ancestry coefficients and optimum K value (K=7) are shown as inferred by rmaverick. Each vertical bar represents an individual haplotype and its membership to the eight cluster are defined by the different colors. Black borders separate the study sites. Samples are ordered chronologically.

**Figure 3.**
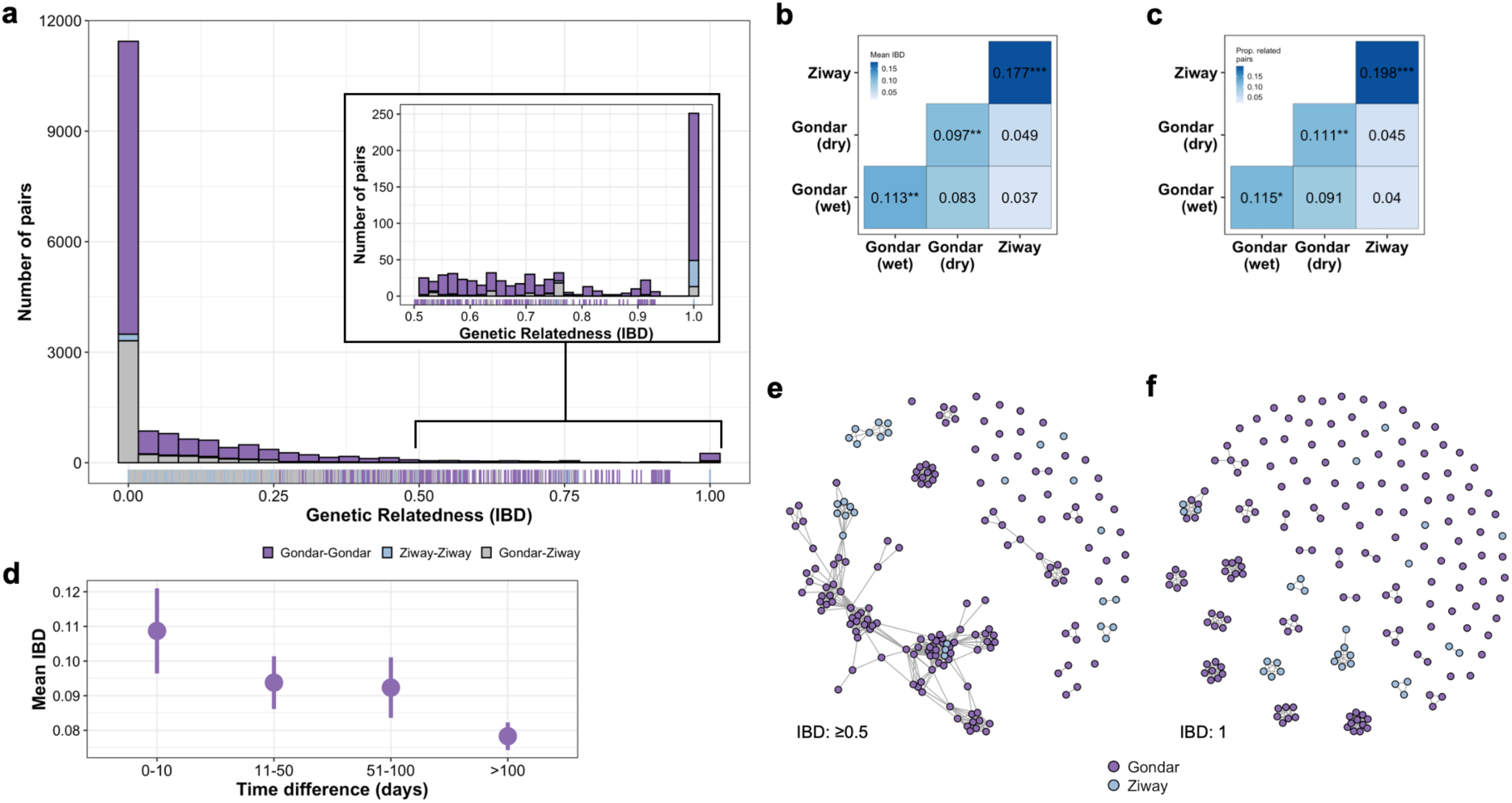
Relatedness between parasites of Ethiopia in Gondar and Ziway (2019–2020). **A**, Histogram of pairwise IBD. Pairwise IBD between all samples (n=17,391), estimated by dcifer (51). Inset shows the heavy tail of the distribution, with some pairs of samples having IBD = 1. **B**, Mean IBD within and between study sites. Asterisks correspond to the permutation test’s p-value, p < 0.05 (*), p < 0.01 (**), and p < 0.001 (***). **C**, Proportion of related infections between study sites. Asterisks correspond to the permutation test’s p-value, p < 0.05 (*), p < 0.01 (**), and p < 0.001 (***). **D,** Temporal patterns of IBD in Gondar. Mean IBD of sample pairs collected 0-10 days apart, 11-50 days apart, 51-100 days apart or >100 days apart. Vertical lines indicate 95% confidence intervals. **E**&**F,** Network of all samples collected from Gondar or Ziway. Edges connecting nodes correspond to (**E**) IBD≥0.5 and (**F**) IBD=1.

**Table 1.**
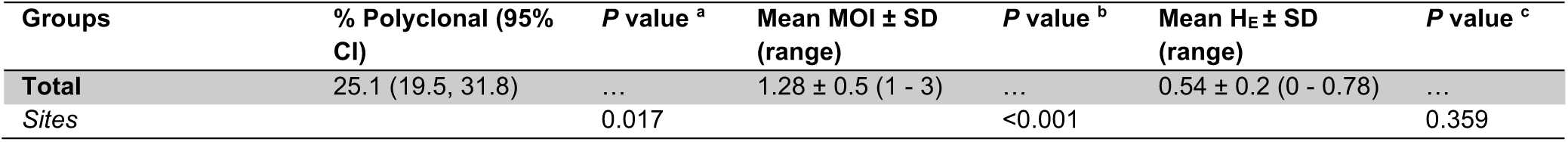
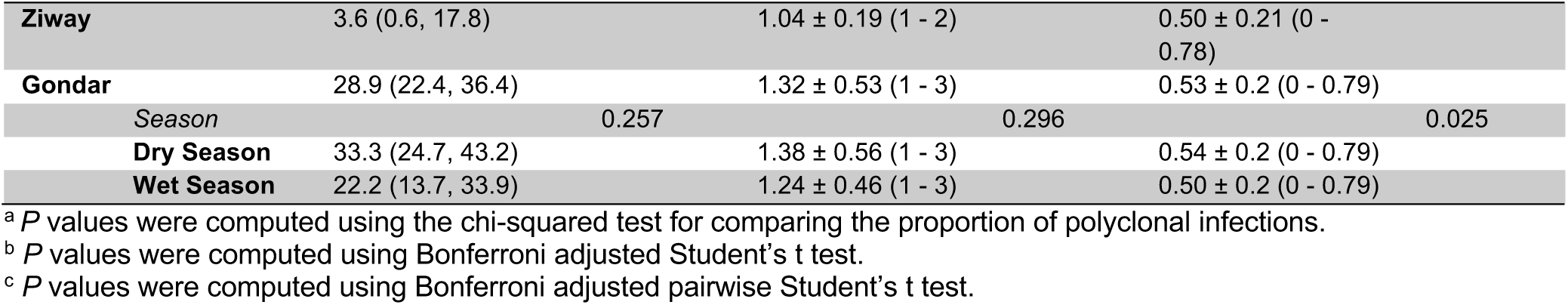
Metrics of Within-Host Diversity and Population-Level genetic diversity in Gondar and Ziway.

### Population genetic structure across Gondar and Ziway

DAPC using the first 31 principal components (PCs), as determined by alpha score optimization, explaining 82.2% of the variation in the original PCA (Supplementary Figure 4a), largely separated the Gondar and Ziway isolates into two distinct clusters (Figure 2c). Population structure analysis by *rmaverick* inferred seven subpopulations (*K*=7) with the highest model evidence (Supplementary Figure 5). We identified some differences between the Gondar and Ziway populations, with cluster 6 found almost exclusively in Ziway (Figure 2d). The separation between Ziway and Gondar was also evident at lower *K* values (Supplementary Figure 5). Inbreeding as determined by mLD was moderate in both study sites but slightly higher in Ziway (Supplementary Table 7).

Among the parasite population in Gondar, we observed subtle separation between the dry and wet seasons by DAPC. No clustering by month or travel history was identified by PCA (Supplementary Figure 4b,c). Population structure analysis revealed no temporal differences between seasons, except for cluster 7, which was found almost exclusively in the wet season, providing evidence that transmission chains were maintained across seasons (Figure 2d).

### Pairwise genetic relatedness between infections

We calculated identity-by-descent (IBD) among all infection pairs (n=17,391) using *dcifer* (51). Expected sharing for full siblings is 0.50, and 0.25 for half-siblings with unrelated parents (60). Among all pairs of parasites, mean IBD was 0.082. We identified a tail of very highly related samples in both sites, suggesting the presence of local clonal transmission chains (Figure 3a, Supplementary Figure 6a,b). In total, we found 1,555 significantly related pairs (8.94%). Mean IBD and the proportion of related infections were higher in Ziway (mean IBD: 0.177; prop. related: 19.8% (75/378)) than in Gondar (mean IBD: 0.093; prop. related: 10.3% (1,288/12,561)) (Figure 3b,c). 1418 (8.15%) pairs were related at the level of half-siblings or higher (IBD ≥0.25), with a higher proportion in Ziway (16.7%) than in Gondar (9.6%). 688 infection pairs (3.96%) were found at the level of full siblings (IBD ≥0.5; Figure 3e), again with a higher proportion in Ziway (11.6%) than Gondar (4.6%). Between the two sites, we observed 4.47% (199/4,452) of related infections at the level of half-siblings or higher and 1.46% (65/4,452) at the level of full siblings or higher.

In Gondar, mean IBD (permutation test, *p*=0.0006) and the proportion of related infections (permutation test, *p*=0.0008) were higher within the same transmission season than between transmission seasons (Figure 3b,c), yet 9.1% of infections were related between seasons, highlighting the overall high relatedness in the study site. Pairs collected temporally closer together showed higher mean IBD than samples collected further apart (Figure 3d).

Among 110 samples (1.44%; 87 from Gondar, 23 from Ziway), 251 pairs were clonal (IBD = 1), with 13 pairs spanning Gondar and Ziway (Figure 3f). Relatedness across a range of IBD thresholds is shown in Supplementary Figure 7, highlighting high relatedness. For example, at IBD ≥0.75, 356 pairs across 125 samples formed 23 clusters with ≥2 samples (Supplementary Figure 7).

In Gondar, the 87 samples with clonal relationships formed 20 different clusters containing two to eleven samples (Figure 4a,b). Ten clusters contained only samples from the same transmission season, and ten clusters contained samples from both transmission seasons. Twelve clusters contained samples collected within <6 months, and eight clusters consisted of samples collected >6 months apart (clusters A, C, E, F, G, I, M, and O), indicating persistent local transmission. The relatedness between clusters was variable, with mean IBD between clusters ranging from 0 to 0.905 (Figure 4c). Of note, in six different clusters, one individual reported recent travel. In all clonal clusters, infections from travelers were the earliest observed cases, with all travelers reporting their infections in November or December 2019. Clonal pairs were found more frequently within the same transmission season than between the two transmission seasons (152 vs. 50, respectively, chi-square test, *p*=0.0012). However, 24.8% (50/202) of clonal pairs were found between transmission seasons, suggesting persistence of clones.

**Figure 4.**
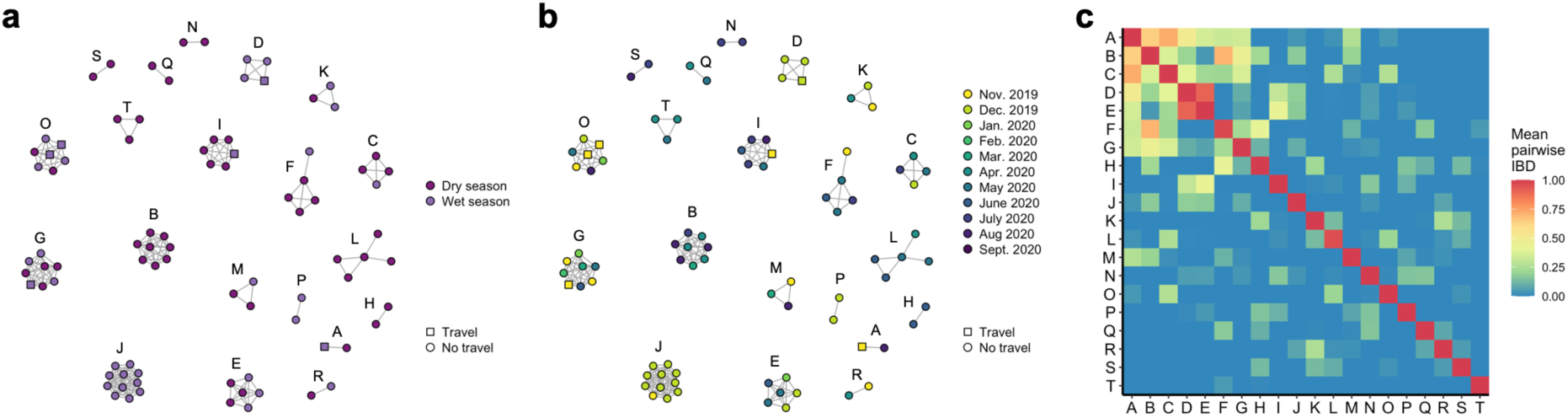
Network of 87 samples collected from Gondar between 2019-2020. **A,** colored by transmission season. **B,** colored by months. Travel (square) and no travel (circle) of individuals are indicated. Edges connecting nodes correspond to IBD=1. **C,** Mean IBD within and between clusters. Mean IBD between clusters ranges from 0 to 0.905 and for within clusters from 0.956 to 1.

### Pfhrp2/3 deletion status

154 samples (134 Gondar, 20 Ziway) with at least 50% sequencing coverage were successfully typed for *pfhrp2*/*3* deletion by ddPCR (Supplementary Figure 8a,d). No associations were found between pfhrp2/3 ddPCR results and parasitemia or total number of reads (Supplementary Figure 8b,c,e,f). 48/154 (31.1%) samples carried *pfhrp2* deletions, and 130/154 (84.4%) carried *pfhrp3* deletions (including mixed infections; Figure 5a,b). *Pfhrp2* deletions were observed only in Gondar. Of 150 samples with data for both genes, 43 (28.7%; including mixed infections) were missing both *pfhrp2* and *pfhrp3*. 2/150 (1.3%) samples carried only the *pfhrp2* deletion and 77/150 (51.3%) samples carried only the *pfhrp3* deletion. 21 (14.0%) samples were wildtype.

**Figure 5.**
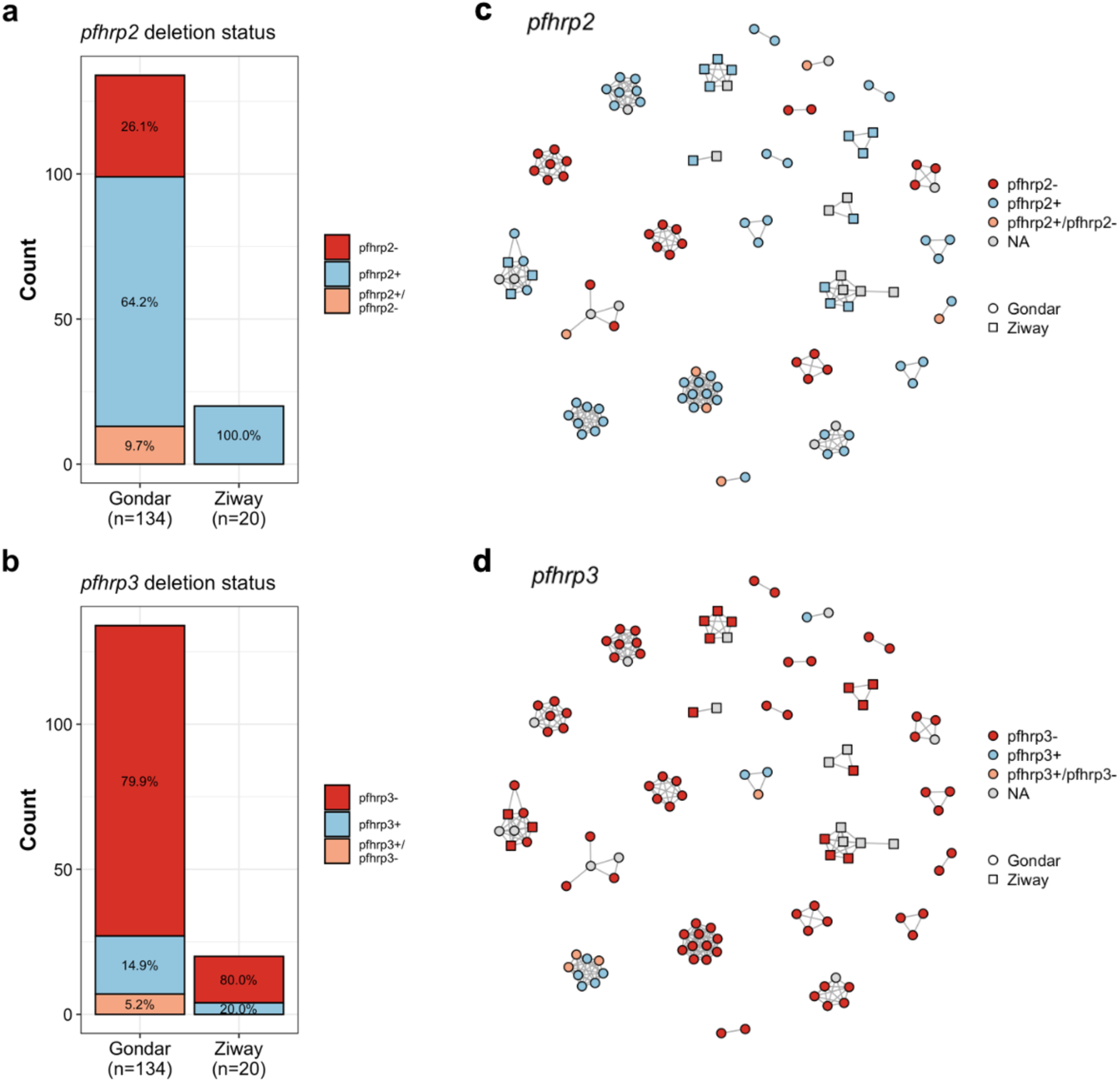
Pfhrp2/3 deletion status by ddPCR. **A,** Pfhrp2 deletion status in 154 samples. Percentage within each study site are indicated. **B,** Pfhrp3 deletion status in 154 samples. Percentage within each study site are indicated. Only samples with at least 10 copies of tRNA were considered for pfhrp2/3 deletion status. **C&D,** Relatedness network of all infections with clonal relationship. Edges connecting nodes correspond to IBD=1. Circles (Gondar) and squared (Ziway) indicate study sites. Colored by **C,** pfhrp2 deletion status and **D,** pfhrp3 deletion status. No clusters show signs of de novo pfhrp2/3 deletions.

Pairwise genetic relatedness was highest among *pfhrp2-*deleted parasites*. Pfhrp2-*deleted parasites showed significantly higher mean IBD, and a four-fold higher proportion of related infections compared to *pfhrp2+* samples (Table 2). No differences in mean IBD and the proportion of related infections were observed between *pfhrp3-* and *pfhrp3+* infections (Table 2). Similarly, estimates of *G*^′′^ between *pfhrp2/3*-deleted parasites and non-deleted controls revealed significant genetic differentiation between *pfhrp2*-deleted and *pfhrp2* wildtype parasites (*G*^′′^ = 0.224, *p*<0.0001), and to a lesser extent, between *pfhrp3*-deleted and non-deleted parasites (*G*^′′^ = 0.097, *p*<0.0001).

**Table 2.** Mean IBD and proportion of related infection pairs between pfhrp2/3-deleted and pfhrp2/3 wildtype parasites. P values of each comparison were calculated by one-sided permutation test (100,000 permutations).

|  | Mean IBD | *P* value | Proportion related infections | *P* value |
| --- | --- | --- | --- | --- |
| ***pfhrp2* deletions** |  | 0.048 |  | <0.00001 |
| ***pfhrp2*<sup>-</sup> (mutant)** | 0.256 |  | 0.362 |  |
| ***pfhrp2*<sup>+</sup> (wildtype)** | 0.082 |  | 0.094 |  |
| ***pfhrp3* deletions** |  | 0.566 |  | 0.898 |
| ***pfhrp3*<sup>-</sup> (mutant)** | 0.091 |  | 0.106 |  |
| ***pfhrp3*<sup>+</sup> (wildtype)** | 0.092 |  | 0.135 |  |

*Pfhrp2* deletions were observed in 10/25 clonal clusters (IBD=1), and *pfhrp3* deletions in 24/25 clusters (including mixed infections; Figure 5c&d). All samples within each clonal cluster were either wildtype or carried the deletion, although some had “mixed” deletion status as expected in polyclonal infections (i.e., *pfhrp2+/pfhrp2-* or *pfhrp3+/pfhrp3-*). No cluster contained both *pfhrp2/3+* and *pfhrp2/3-* clones. At a sub-clonal relatedness threshold, e.g., at IBD >0.5, samples with and without deletions were observed in the same cluster, reflective of Mendelian inheritance of the deletion (Supplementary Figures 9 and 10).

### Prevalence of drug-resistance markers

Data for five drug resistance markers (*pfdhfr*, *pfdhps*, *pfmdr1*, *pfmdr2*, and *pfk13*) were obtained for 94% to 98% of samples per marker (Table 3). Frequencies of mutant alleles were high in all genes, except for *pfk13*, where no mutations were found. Mutations associated with sulfadoxine-pyrimethamine (SP) resistance in both the *pfdhfr* (N51I, C59R and S108N) and *pfdhps* (K540E) genes were observed at frequencies of up to >90%. Over 97% of infections carried the *pfmdr1* N86 wildtype and 184F mutant alleles, both of which are associated with decreased sensitivity to lumefantrine (61,62). A quarter of all infections carried the *pfmdr2* I492V mutation, which has been implicated in artemisinin resistance (63,64). Mutation frequencies for *pfdhfr* C59R and *pfmdr2* I492V differed significantly between Gondar and Ziway (Supplementary Table 8)

**Table 3.** Frequency of mutations of drug resistance alleles from 187 samples from Gondar (n=159) and Ziway (n=28).

| Gene | Chromosome | Position | Mutation | REF | ALT | No. successfully sequenced (%) | Frequency (%)* |  |  |
| --- | --- | --- | --- | --- | --- | --- | --- | --- | --- |
|  |  |  |  |  |  |  | Wild-type | Mutant | Mixed |
| *pfdhfr* | chr4 | 748239 | N51I | A | T | 177 (94.7%) | 0 | 100 | 0 |
|  | chr4 | 748262 | C59R | T | C | 177 (94.7%) | 54.8 | 42.9 | 2.3 |
|  | chr4 | 748410 | S108N | G | A | 177 (94.7%) | 0.6 | 97.7 | 1.7 |
|  | chr4 | 748410 | S108T | G | C | 177 (94.7%) | 100 | 0 | 0 |
|  | chr4 | 748577 | I164L | A | T | 177 (94.7%) | 100 | 0 | 0 |
| *pfdhps* | chr8 | 549993 | K540E | A | G | 177 (94.7%) | 7.3 | 92.7 | 0 |
|  | chr8 | 550117 | A581G | C | G | 177 (94.7%) | 100 | 0 | 0 |
|  | chr8 | 550212 | A613T | G | A | 177 (94.7%) | 100 | 0 | 0 |
|  | chr8 | 550212 | A613S | G | T | 177 (94.7%) | 100 | 0 | 0 |
| *pfmdr1* | chr5 | 958145 | N86Y | A | T | 178 (95.2%) | 97.2 | 2.8 | 0 |
|  | chr5 | 958440 | Y184F | A | T | 184 (98.4%) | 1.1 | 97.3 | 1.6 |
| *pfmdr2* | chr14 | 1956202 | I492V | A | G | 176 (94.1%) | 64.2 | 26.1 | 9.7 |
|  | chr14 | 1956225 | T484I | C | T | 176 (94.1%) | 100 | 0 | 0 |
| *pfk13* | chr13 | 1725370 | I543T | A | G | 182 (97.3%) | 100 | 0 | 0 |
|  | chr13 | 1725382 | R539T | C | G | 182 (97.3%) | 100 | 0 | 0 |
|  | chr13 | 1725385 | G538V | G | T | 182 (97.3%) | 100 | 0 | 0 |
|  | chr13 | 1725388 | N537I | A | T | 182 (97.3%) | 100 | 0 | 0 |
|  | chr13 | 1725418 | P527H | C | A | 182 (97.3%) | 100 | 0 | 0 |
|  | chr13 | 1725521 | Y493H | T | C | 182 (97.3%) | 100 | 0 | 0 |
|  | chr13 | 1725556 | A481V | C | T | 182 (97.3%) | 100 | 0 | 0 |

\* Polyclonal and monoclonal samples were included. "Mixed" indicates infections with both the mutant and the wild-type allele.

In Gondar, all of the 97 monoclonal samples with complete allele calls for all drug resistance loci carried resistance-associated alleles in at least three of the four genes (Figure 6a). The common *pfdhfr* triple mutant (51I-59R-108N) was observed in 39/97 (40.2%) isolates, and 36/97 (37.1%) also carried the *pfdhps* 540E mutation. All but one isolates carried the *pfdhfr* 51I and 108N mutations. We observed 93/97 (95.9%) isolates carrying the *pfmdr1* N86 wildtype and 184F mutant alleles. In Ziway, all isolates carried the *pfdhfr* triple mutant alleles (51I-59R-108N), the *pfdhps* 540E mutant allele, and the *pfmdr1* N86 wild type and 184F mutant alleles (Figure 6b).

**Figure 6.**
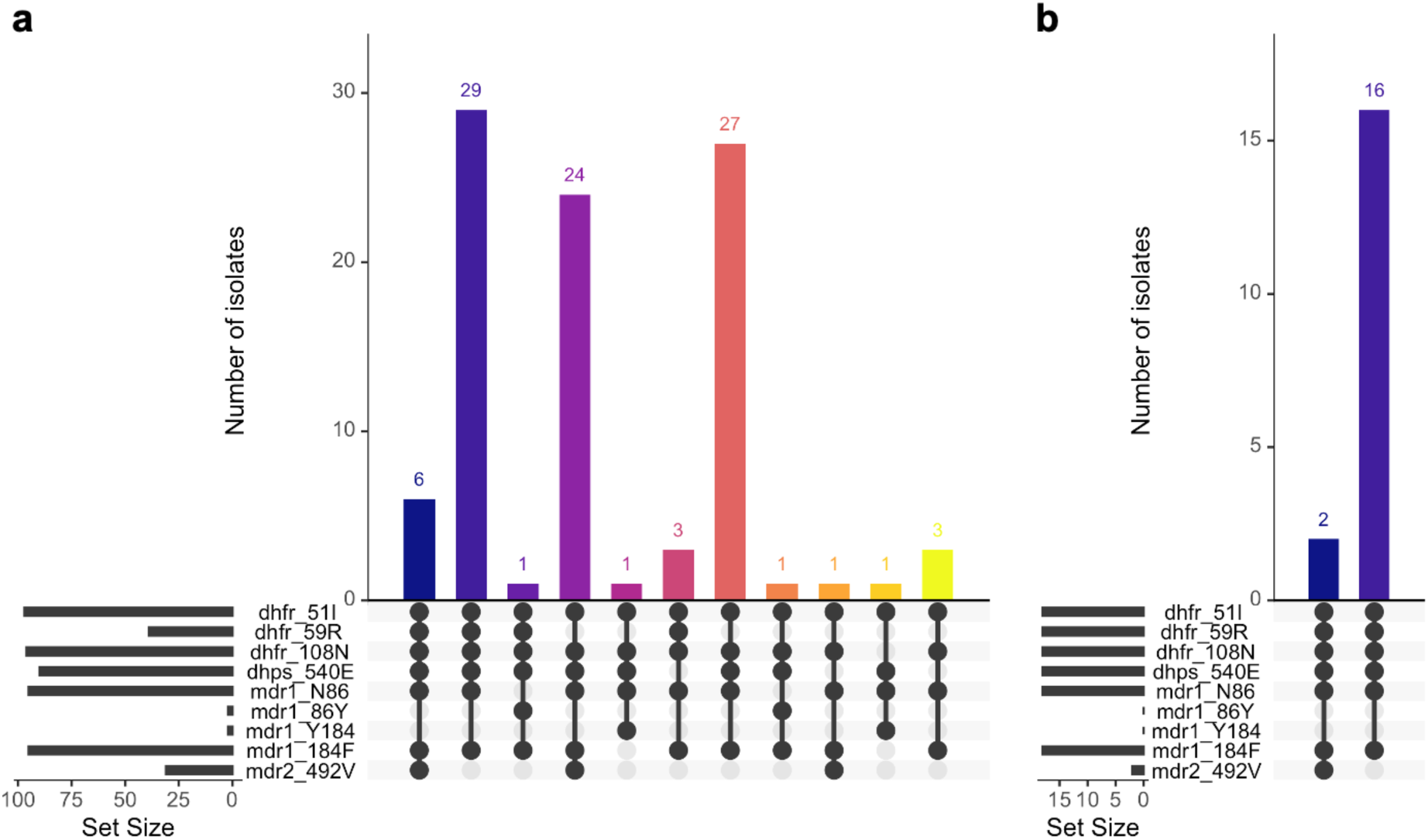
UpSetR plot showing the number of isolates from monoclonal samples. **A**, Samples from Gondar with complete allele calls (n=97) carrying different combinations of antimalarial drug resistance-associated alleles in four genes (pfdhfr, pfdhps, pfmdr1, and pfmdr2). **B**, Samples from Ziway with complete allele calls (n=18).

### Genetic relatedness to samples from Africa and Asia

To investigate connectivity within Ethiopia, we estimated pairwise genetic relatedness by IBD between samples from our study (n=187) and published genomes from Ethiopia (n=21, MalariaGEN), collected in 2013 and 2015, respectively. Mean IBD between all pairs of infection from our study and publicly available samples from Ethiopia was 0.089. We found 11.71% (460/3,927) significantly related pairs between samples from our study and other samples from Ethiopia. All 21 MalariaGEN samples were significantly related to at least one sample from our study, indicating a high degree of relatedness of infections within Ethiopia. We identified 9.32% (366/3,927) pairs at the relatedness level of half-siblings (IBD ≥0.25), 1.66% (65/3,927) at the level of full-siblings (IBD ≥0.5), and 0.36% (14/3,927) highly related pairs with an IBD ≥0.9. Relatedness networks at different thresholds are shown in Supplementary Figure 11.

A previous major population genomic study in Africa found that Ethiopia has a distinctive population of *P. falciparum* compared to other regions, suggesting a distinct ancestry from the rest of Africa (23). We performed DAPC on samples from our study and publicly available genomes from 10 countries from Africa (n=140) and 6 countries from Southeast Asia (n=58) (Supplementary Figure 12) to investigate the genetic differentiation between our samples and other isolates across the world. Consistent with previous findings, samples from our study sites formed a separate cluster together with other Ethiopian samples (Figure 7a). The remaining isolates from Africa formed another cluster, as did all isolates from Southeast Asia. Relatedness analysis by IBD confirmed that samples from our two study sites were most closely related to other samples from Ethiopia (Figure 7b,c, Supplementary Tables 9-11). Relatedness to other African countries, even nearby East African countries, and countries in Southeast Asia was very low, confirming previous findings that Ethiopia has a distinct *P. falciparum* population (23).

**Figure 7.**
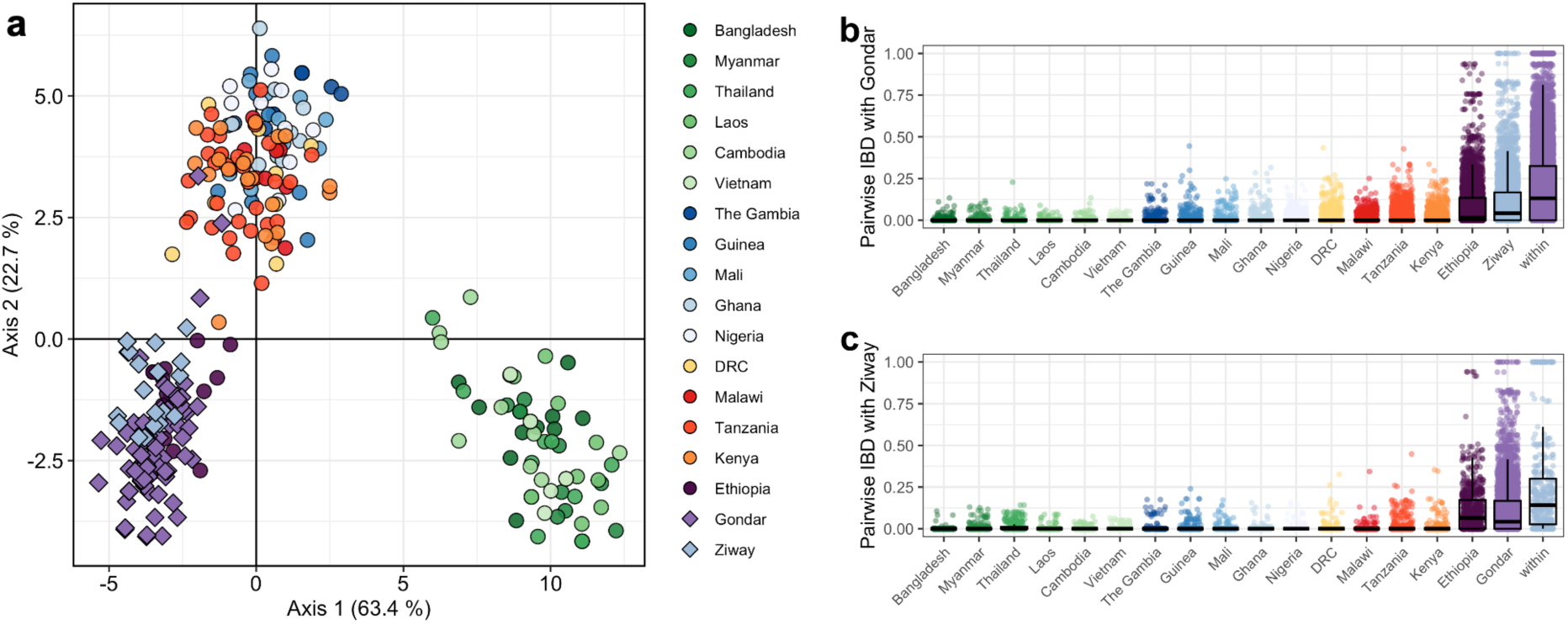
**A,** Genetic differentiation between global P. falciparum populations by discriminatory analysis of principal components (DAPC) using 28 microhaplotypes. The first 24 components of a principal component analysis (PCA) are shown. Each point represents a single isolate (n=338); colors indicate the country of origin. Only monoclonal samples were used for this analysis (n=113 Gondar; n=27 Ziway). **B,** Boxplot showing pairwise IBD between samples from Gondar (n=159) and several countries from Africa (n=140) and Southeast Asia (n=58). C, Boxplot showing pairwise IBD between Ziway (n=28) and several countries from Africa (n=140) and Southeast Asia (n=58).

## DISCUSSION

The *P. falciparum* population in Gondar in the Ethiopian highlands exhibits moderate genetic diversity and low multiplicity, in line with studies from other sites in Ethiopia (23–25). A high proportion of closely related infections was found, and virtually all infections linked at the level of half-siblings (IBD ≥0.25) or higher. Historically, areas above 2,000 m were considered malaria-free in Ethiopia (65), but in Gondar region, high number of cases were observed during the wet season (Ewnetu, et al., unpublished data). Seasonal migration from highly endemic lowland areas may contribute significantly to highland malaria transmission (8–10), but the dynamics of highland transmission remain elusive.

Where transmission is low, the frequency of mixed genotypes decreases, leading to self-fertilization and increased genetic relatedness (60,66,67). In contrast, imported infections are expected to show low levels of relatedness. A high degree of genetic relatedness (IBD) among infections within Gondar and Ziway revealed patterns of ongoing local transmission. In Gondar, MOI and genetic diversity were slightly higher in dry season infections. This pattern may indicate that importation plays a greater role in the dry season, while clones are transmitted locally in the wet season, resulting in clusters of closely related infections. Yet, several clonal clusters consisted of samples from both transmission seasons collected >6 months apart, providing evidence for ongoing local transmission.

Five clonal clusters included samples with travel history, and all samples from travelers were collected at the beginning of the study in November and December 2019. These infections could potentially reflect imported cases from returning seasonal migrants to Gondar, which then lead to ongoing local transmission. However, only a small proportion of patients reported travel, so parasite importation is unlikely to be the main driver of persistent malaria transmission in the highlands around Gondar. For the current study, only symptomatic and microscopy-positive samples were sequenced. Travelers and individuals with secondary infections may remain asymptomatic. Future studies could include submicroscopic and asymptomatic carriers, who are likely to contribute substantially to transmission in Ethiopia (43,68,69), to fully understand the impact of imported infections.

The Horn of Africa is heavily affected by *pfhrp2/3*-deleted parasites, as evidenced by reports from Eritrea (34,35), Djibouti (36,37), Sudan (38), South Sudan (38), and also Ethiopia (31–33). *Pfhrp2* deletions have recently been described and estimated at 11.5% in parts of the Amhara region (32). The higher overall frequency of *pfhrp3*-deleted parasites compared to *pfhrp2* deletions observed in Gondar and Ziway corroborates previous findings in different geographical locations (30,52). A recent study from Ethiopia suggests that the *pfhrp2* deletion mutation emerged and recently expanded from a single origin, whereas *pfhrp3* deletion mutations expanded in the more distant past and likely have multiple independent origins (32). Similarly, a study from Eritrea found that 30 of 31 (96.8%) *pfhrp2-*deleted strains fell into a single genetically related cluster (34). Our data support these findings. *Pfhrp2-* samples had a significantly higher mean IBD compared to *pfhrp2+* samples, had a fourfold higher proportion of related infections, and the relatedness between *pfhrp2-* and *pfhrp2+* samples was low. In contrast, no differences in relatedness were observed by *pfhrp3* status. Mapping of clonal clusters (IBD = 1) suggests that frequent *de novo* mutation is not a major force structuring *pfhrp2/3* deletion. Analysis of genetic differentiation (i.e., *G*^′′^) confirmed that *pfhrp2*-deleted parasites were genetically distinct from wildtype parasites, and to a lesser extent also for *pfhrp3*.

Drug resistance is a major threat to malaria control. Since 2004, the first-line treatment in Ethiopia for *P. falciparum* has been artemether-lumefantrine (AL) (40). Previous studies have reported variable prevalence of mutations in the drug-resistance loci *pfdhps*, *pfdhfr, and pfmdr1* across Ethiopia (25–29). Consistent with previous reports from different parts of Ethiopia (25,26), mutations associated with sulfadoxine-pyrimethamine (SP) resistance in both *pfdhfr* (N51I and S108N) and *pfdhps* (K540E) genes were observed at near-fixation in Gondar and Ziway. Other mutations were common, including in *pfmdr1*, which are associated with decreased sensitivity to lumefantrine (61,62). These findings warrant monitoring of the efficacy of AL as a first-line treatment for *P. falciparum* in Ethiopia and may require a re-evaluation of AL. In addition, one-quarter of all infections harbored the *pfmdr2* 492V mutant allele, which has been implicated in partial artemisinin resistance (63,64). However, our study did not identify any validated *pfk13* artemisinin partial resistance-conferring mutations.

Consistent with previous results (23), our samples, along with publicly available Ethiopian samples, formed a distinct cluster, separated from all other African isolates. The separation from East African populations is noteworthy because, despite the increasing mobility of people between Ethiopia and the rest of Africa, particularly East Africa, there still appear to be significant barriers to gene flow. These barriers may also explain why *pfhrp2/3* deletions appear to be disproportionately prevalent in the Horn of Africa (31–38). The genetic diversity of *P. falciparum* infections in Gondar and Ziway was markedly lower (mean H_E_: 0.54) than previously observed in another low transmission area in Zanzibar (mean H_E_: 0.73), using the same microhaplotype panel (39), suggesting a smaller parasite population size in Gondar. The direct comparison between two elimination sites corroborates the usefulness of malaria genomic surveillance to understand transmission intensities in different sites.

In conclusion, we show a predominance of low-complexity infections and a significant percentage of shared genomic haplotypes in the population, providing evidence for persistent local and focal transmission. However, we also identify potential imported cases contributing to onward transmission. We corroborate previous data on *pfhrp2/3* deletion and drug resistance, both of which pose a threat to malaria control in Ethiopia. We demonstrate the utility of genetic tools to inform interventions to eliminate local transmission and prevent further introduction of imported malaria parasites that may contribute to local transmission.

## DATA AVAILABILITY

All raw sequencing data is available at the NCBI SRA (Accession number: PRJNA962572). De-identified datasets generated during the current study and used to make all figures are available as supplementary files or tables. This publication uses data from the MalariaGEN *Plasmodium falciparum* Community Project as described in ‘Pf7: an open dataset of *Plasmodium falciparum* genome variation in 20,000 worldwide samples’, MalariaGEN *et al* (53).

## CODE AVAILABILITY

All analyses were completed in R version 4.1.2 using published R packages, as described in the methods section. This study did not generate any novel code. Code for analyses and raw analysis results are available upon request.

## Notes

### Competing Interest Statement

The authors have declared no competing interest.

