## Supplementary Information for "*Plasmodium falciparum* transmission in the highlands of Ethiopia is driven by closely related and clonal parasites"

**TABLE OF CONTENTS**


| | |
|---|---|
| 1. Supplementary Table 1 | Page 2 |
| 2. Supplementary Table 2 | Page 3 |
| 3. Supplementary Table 3 | Page 5 |
| 4. Supplementary Table 4 | Page 5 |
| 5. Supplementary Table 5 | Page 6 |
| 6. Supplementary Table 6 | Page 7 |
| 7. Supplementary Table 7 | Page 11 |
| 8. Supplementary Table 8 | Page 12 |
| 9. Supplementary Table 9 | Page 13 |
| 10. Supplementary Table 10 | Page 14 |
| 11. Supplementary Table 11 | Page 15 |
| 12. Supplementary Figure 1 | Page 16 |
| 13. Supplementary Figure 2 | Page 17 |
| 14. Supplementary Figure 3 | Page 18 |
| 15. Supplementary Figure 4 | Page 19 |
| 16. Supplementary Figure 5 | Page 20 |
| 17. Supplementary Figure 6 | Page 21 |
| 18. Supplementary Figure 7 | Page 22 |
| 19. Supplementary Figure 8 | Page 23 |
| 20. Supplementary Figure 9 | Page 24 |
| 21. Supplementary Figure 10 | Page 25 |
| 22. Supplementary Figure 11 | Page 26 |
| 23. Supplementary Figure 12 | Page 27 |

### SUPPLEMENTARY TABLES

***Supplementary Table 1**. Overview and genomic location of the 35 loci included in the panel.*

| name | seqnames | start | end | width | type | geneID | description |
|---|---|---|---|---|---|---|---|
| t01 | Pf3D7_01_v3 | 145417 | 145655 | 239 | Microhaplotype | PF3D7_0103300 | conserved protein, unknown function |
| cpmp_22 | Pf3D7_01_v3 | 180130 | 180370 | 241 | Microhaplotype | PF3D7_0104100 | conserved Plasmodium membrane protein, unknown function |
| t04 | Pf3D7_01_v3 | 495942 | 496171 | 230 | Microhaplotype | PF3D7_0113100 | surface-associated interspersed protein 1.1 (SURFIN 1.1) |
| t08 | Pf3D7_01_v3 | 534183 | 534397 | 215 | Microhaplotype | PF3D7_0113800 | DBL containing protein, unknown function |
| t11 | Pf3D7_02_v3 | 278145 | 278369 | 225 | Microhaplotype | PF3D7_0207000 | merozoite surface protein 4 |
| csp_27 | Pf3D7_03_v3 | 221468 | 221655 | 188 | Microhaplotype | PF3D7_0304600 | circumsporozoite (CS) protein |
| t17 | Pf3D7_03_v3 | 618368 | 618613 | 246 | Microhaplotype | PF3D7_0315200 | circumsporozoite- and TRAP-related protein |
| t25 | Pf3D7_04_v3 | 748201 | 748469 | 269 | Drug Resistance | PF3D7_0417200 | bifunctional dihydrofolate reductase-thymidylate synthase |
| t26 | Pf3D7_04_v3 | 748501 | 748725 | 225 | Drug Resistance | PF3D7_0417200 | bifunctional dihydrofolate reductase-thymidylate synthase |
| t28 | Pf3D7_04_v3 | 1037603 | 1037868 | 266 | Microhaplotype | PF3D7_0422500 | pre-mRNA-splicing helicase BRR2, putative |
| t30 | Pf3D7_04_v3 | 1102363 | 1102606 | 244 | Microhaplotype | PF3D7_0424400 | surface-associated interspersed protein 4.2 (SURFIN 4.2) |
| t31 | Pf3D7_04_v3 | 1113423 | 1113632 | 210 | Microhaplotype | PF3D7_0424600 | Plasmodium exported protein (PHISTb), unknown function |
| t33 | Pf3D7_05_v3 | 329352 | 329578 | 227 | Microhaplotype | PF3D7_0508000 | 6-cysteine protein |
| t34 | Pf3D7_05_v3 | 958028 | 958254 | 227 | Drug Resistance | PF3D7_0523000 | multidrug resistance protein 1 |
| t35 | Pf3D7_05_v3 | 958363 | 958539 | 177 | Drug Resistance | PF3D7_0523000 | multidrug resistance protein 1 |
| t45 | Pf3D7_07_v3 | 1358667 | 1358941 | 275 | Microhaplotype | PF3D7_0731500 | erythrocyte binding antigen-175 |
| t47 | Pf3D7_08_v3 | 336439 | 336674 | 236 | Microhaplotype | PF3D7_0806200 | Dpy-19-like C-mannosyltransferase, putative |
| t49 | Pf3D7_08_v3 | 549963 | 550251 | 289 | Drug Resistance | PF3D7_0810800 | hydroxymethyldihydropterin pyrophosphokinase-dihydropteroate synthase |
| t53 | Pf3D7_08_v3 | 1362864 | 1363120 | 257 | Microhaplotype | PF3D7_0831600 | cytoadherence linked asexual protein 8 |
| t54 | Pf3D7_09_v3 | 516905 | 517125 | 221 | Microhaplotype | PF3D7_0911300 | cysteine repeat modular protein 1 |
| t60 | Pf3D7_10_v3 | 377063 | 377237 | 175 | Microhaplotype | PF3D7_1009200 | ribonuclease, putative |
| t62 | Pf3D7_10_v3 | 1386673 | 1386902 | 230 | Microhaplotype | PF3D7_1035000 | U2 snRNA/tRNA pseudouridine synthase, putative |
| t66 | Pf3D7_11_v3 | 1009832 | 1010071 | 240 | Microhaplotype | PF3D7_1125800 | kelch domain-containing protein, putative |
| t67 | Pf3D7_11_v3 | 1018921 | 1019113 | 193 | Microhaplotype | PF3D7_1126100 | autophagy-related protein 7, putative |
| ama1_D2_18 | Pf3D7_11_v3 | 1294271 | 1294520 | 250 | Microhaplotype | PF3D7_1133400 | apical membrane antigen 1 |
| t72 | Pf3D7_12_v3 | 63136 | 63313 | 178 | Microhaplotype | PF3D7_1200700 | acyl-CoA synthetase |
| t73 | Pf3D7_12_v3 | 659859 | 660037 | 179 | Microhaplotype | PF3D7_1216600 | cell traversal protein for ookinetes and sporozoites |
| t74 | Pf3D7_12_v3 | 684060 | 684289 | 230 | Microhaplotype | PF3D7_1217300 | GTP-binding protein EngA |
| msp7_1 | Pf3D7_13_v3 | 1419369 | 1419544 | 176 | Microhaplotype | PF3D7_1335100 | merozoite surface protein 7 |
| t82 | Pf3D7_13_v3 | 1725338 | 1725592 | 255 | Drug Resistance | PF3D7_1343700 | kelch protein K13 |
| t85 | Pf3D7_13_v3 | 2124603 | 2124876 | 274 | Microhaplotype | PF3D7_1353100 | Plasmodium exported protein, unknown function |
| t96 | Pf3D7_14_v3 | 1956097 | 1956312 | 216 | Drug Resistance | PF3D7_1447900 | multidrug resistance protein 2 |
| t98 | Pf3D7_14_v3 | 2524932 | 2525119 | 188 | Microhaplotype | PF3D7_1462300 | GTP-binding protein, putative |
| cpp_30 | Pf3D7_14_v3 | 3121036 | 3121267 | 232 | Microhaplotype | PF3D7_1475800 | conserved Plasmodium protein, unknown function |
| t99 | Pf3D7_14_v3 | 3124615 | 3124870 | 256 | Microhaplotype | PF3D7_1475900 | KELT protein |

***Supplementary Table 2.*** *Primer sequences for the 35-plex panel. Final primer pools are generated by combining 10µL of each primer stock (total 700 µL) with 300 µL NF ddH2O. This is then used to prepare the PCR reaction.*

| **Primer** | **Direction** | **Sequence (5' - 3')** | **Stock concentration (µM)** | **Concentration in final pool (µM)** | **Final concentration in PCR reaction (µM)** |
|---|---|---|---|---|---|
| ama1_D2_18 | Forward | GTGACCTATGAACTCAGGAGTCGAACTCAATATAGACTTCCATCAGG | 1000 | 10 | 0.227 |
| cpmp_22 | Forward | GTGACCTATGAACTCAGGAGTCGGAAGCTATAGGTATCAGATCC | 1000 | 10 | 0.227 |
| cpp_30 | Forward | GTGACCTATGAACTCAGGAGTCAACACAATCTTCCTTAGCCAATTC | 1000 | 10 | 0.227 |
| csp_27 | Forward | GTGACCTATGAACTCAGGAGTCGACCCAAACCGAAATGTAGATG | 1000 | 10 | 0.227 |
| msp7_1 | Forward | GTGACCTATGAACTCAGGAGTCGCACAAGGAGGAGAATCGAC | 1000 | 10 | 0.227 |
| t01 | Forward | GTGACCTATGAACTCAGGAGTCTTCGATATGTTTAAATATATGATTCTCG | 1000 | 10 | 0.227 |
| t11 | Forward | GTGACCTATGAACTCAGGAGTCCTTACCATTCGCGCTTTCTTG | 1000 | 10 | 0.227 |
| t17 | Forward | GTGACCTATGAACTCAGGAGTCGACACTACAGACAAAAATAAATGATC | 1000 | 10 | 0.227 |
| t25 | Forward | GTGACCTATGAACTCAGGAGTCCTAGGAAATAAAGGAGTATTACCATG | 1000 | 10 | 0.227 |
| t26 | Forward | GTGACCTATGAACTCAGGAGTCTGTTTATATCATTAACAAAGTTGAAGATC | 1000 | 10 | 0.227 |
| t28 | Forward | GTGACCTATGAACTCAGGAGTCTTGAAGTACAATATGAAATCGATCTTG | 1000 | 10 | 0.227 |
| t30 | Forward | GTGACCTATGAACTCAGGAGTCTAGAGGTGTTGATGTTAATATGGAG | 1000 | 10 | 0.227 |
| t31 | Forward | GTGACCTATGAACTCAGGAGTCTCTGCATGCAGTAATGAATCTATTG | 1000 | 10 | 0.227 |
| t33 | Forward | GTGACCTATGAACTCAGGAGTCGTTAAAACAACAGGAGGAACACTAA | 1000 | 10 | 0.227 |
| t34 | Forward | GTGACCTATGAACTCAGGAGTCAAATGTTTACCTGCACAACATAGAAA | 1000 | 10 | 0.227 |
| t35 | Forward | GTGACCTATGAACTCAGGAGTCGAACAAGTGAGTTCAGGAATTGG | 1000 | 10 | 0.227 |
| t04 | Forward | GTGACCTATGAACTCAGGAGTCCACCAAAATATTATATACCACAAGAC | 1000 | 10 | 0.227 |
| t45 | Forward | GTGACCTATGAACTCAGGAGTCAAGGATCATTTCATTGAAGCCTCT | 1000 | 10 | 0.227 |
| t47 | Forward | GTGACCTATGAACTCAGGAGTCATACATGAATATGATTAACAACGAACC | 1000 | 10 | 0.227 |
| t49 | Forward | GTGACCTATGAACTCAGGAGTCAAAAGAGGAAATCCACATACAATGG | 1000 | 10 | 0.227 |
| t53 | Forward | GTGACCTATGAACTCAGGAGTCCGACTTCTTTAAGAGAACAAACAG | 1000 | 10 | 0.227 |
| t54 | Forward | GTGACCTATGAACTCAGGAGTCTGTGGGCGCAAAAACTATAAATGA | 1000 | 10 | 0.227 |
| t60 | Forward | GTGACCTATGAACTCAGGAGTCCAAATAGCATTCGATAAATTTGAAAATTTC | 1000 | 10 | 0.227 |
| t62 | Forward | GTGACCTATGAACTCAGGAGTCCTACAGTTATATAGGACGATTTTGG | 1000 | 10 | 0.227 |
| t66 | Forward | GTGACCTATGAACTCAGGAGTCCTGTGAGCATATCTTGGTTCAG | 1000 | 10 | 0.227 |
| t67 | Forward | GTGACCTATGAACTCAGGAGTCTACTACTACTTATGTTACTTATACCAC | 1000 | 10 | 0.227 |
| t72 | Forward | GTGACCTATGAACTCAGGAGTCATTCTATTAAAAACTTTGGGTACTCC | 1000 | 10 | 0.227 |
| t73 | Forward | GTGACCTATGAACTCAGGAGTCCTGGTACTATTATACCATATGTTGC | 1000 | 10 | 0.227 |
| t74 | Forward | GTGACCTATGAACTCAGGAGTCCTACCATTTTTAATCGACTAACTCG | 1000 | 10 | 0.227 |
| t08 | Forward | GTGACCTATGAACTCAGGAGTCCAGATGTGATAAATATATGTGACATTTG | 1000 | 10 | 0.227 |
| t82 | Forward | GTGACCTATGAACTCAGGAGTCGTATAATAGAAGAGCCATCATATCC | 1000 | 10 | 0.227 |
| t85 | Forward | GTGACCTATGAACTCAGGAGTCATTTGGTTTCAATAAAATTATCAGCTTTC | 1000 | 10 | 0.227 |
| t96 | Forward | GTGACCTATGAACTCAGGAGTCTTTTCTCCACTTTGTAATTTTTATTGTTG | 1000 | 10 | 0.227 |
| t98 | Forward | GTGACCTATGAACTCAGGAGTCGATCTTCTCTGAAATTTACTTGAATTG | 1000 | 10 | 0.227 |
| t99 | Forward | GTGACCTATGAACTCAGGAGTCGATTATTTGGTAATGAACAACCAGG | 1000 | 10 | 0.227 |
| ama1_D2_18 | Reverse | CTGAGACTTGCACATCGCAGCCCTGCATGTCTTGAACATAAAGTC | 1000 | 10 | 0.227 |
| cpmp_22 | Reverse | CTGAGACTTGCACATCGCAGCTAGAATACGTGCTTTATAAACAAAGAG | 1000 | 10 | 0.227 |

| | | | | | |
|---|---|---|---|---|---|
| cpp_30 | Reverse | CTGAGACTTGCACATCGCAGCATTACTACCTTTCAGCATATCCGA | 1000 | 10 | 0.227 |
| csp_27 | Reverse | CTGAGACTTGCACATCGCAGCGAGCCAGGCTTTATTCTAACTTG | 1000 | 10 | 0.227 |
| msp7_1 | Reverse | CTGAGACTTGCACATCGCAGCGAATTTAAAACGAAGTCTTCATATTC | 1000 | 10 | 0.227 |
| t01 | Reverse | CTGAGACTTGCACATCGCAGCTGTCAAGGTATATTAAAGTATGGTATC | 1000 | 10 | 0.227 |
| t11 | Reverse | CTGAGACTTGCACATCGCAGCAATATAGTACCAGAAAATGGAAGAATG | 1000 | 10 | 0.227 |
| t17 | Reverse | CTGAGACTTGCACATCGCAGCGAAACTCCTACTACCAATAATTTGAC | 1000 | 10 | 0.227 |
| t25 | Reverse | CTGAGACTTGCACATCGCAGCAATATAACATTTATCCTATTGCTTAAAGG | 1000 | 10 | 0.227 |
| t26 | Reverse | CTGAGACTTGCACATCGCAGCACATCGCTAACAGAAATAATTTGATAC | 1000 | 10 | 0.227 |
| t28 | Reverse | CTGAGACTTGCACATCGCAGCGTATTTGGTTTGTCAGGCAATTCG | 1000 | 10 | 0.227 |
| t30 | Reverse | CTGAGACTTGCACATCGCAGCCCAAACATACATCCTCTTTTTGTTC | 1000 | 10 | 0.227 |
| t31 | Reverse | CTGAGACTTGCACATCGCAGCGTAGCATGCTCAGATATTATGATG | 1000 | 10 | 0.227 |
| t33 | Reverse | CTGAGACTTGCACATCGCAGCTTGTAGAATATTCGTGTCCGTATAG | 1000 | 10 | 0.227 |
| t34 | Reverse | CTGAGACTTGCACATCGCAGCGATGTAATTACATCCATACAATAACTTG | 1000 | 10 | 0.227 |
| t35 | Reverse | CTGAGACTTGCACATCGCAGCTTTCTTATTACATATGACACCACAAAC | 1000 | 10 | 0.227 |
| t04 | Reverse | CTGAGACTTGCACATCGCAGCGGAAAATCTTTGGTGGGAAAAATAG | 1000 | 10 | 0.227 |
| t45 | Reverse | CTGAGACTTGCACATCGCAGCAAAGTTTCTCTCTAAATTCATTCCAC | 1000 | 10 | 0.227 |
| t47 | Reverse | CTGAGACTTGCACATCGCAGCACTAAAAGGAAGACAAGCTGAAAAC | 1000 | 10 | 0.227 |
| t49 | Reverse | CTGAGACTTGCACATCGCAGCATTTATTACAACATTTTGATCATTCATGC | 1000 | 10 | 0.227 |
| t53 | Reverse | CTGAGACTTGCACATCGCAGCAGTTTTCTTCTGAATAGTTCTACAAAG | 1000 | 10 | 0.227 |
| t54 | Reverse | CTGAGACTTGCACATCGCAGCTTGTATATCCAAAATTACATTGACAATTG | 1000 | 10 | 0.227 |
| t60 | Reverse | CTGAGACTTGCACATCGCAGCCATCTATATAGTCCCCGATGAATG | 1000 | 10 | 0.227 |
| t62 | Reverse | CTGAGACTTGCACATCGCAGCAATGAGGACAAGGAAAATAATACATTC | 1000 | 10 | 0.227 |
| t66 | Reverse | CTGAGACTTGCACATCGCAGCATGAAACATATAAATTTGATTTCCAAGC | 1000 | 10 | 0.227 |
| t67 | Reverse | CTGAGACTTGCACATCGCAGCCTCCTTTAGGTATTACGGTAGC | 1000 | 10 | 0.227 |
| t72 | Reverse | CTGAGACTTGCACATCGCAGCACATTTATCTTATTTACCCGTATCTC | 1000 | 10 | 0.227 |
| t73 | Reverse | CTGAGACTTGCACATCGCAGCTCACCAACCTTTTTAGAATCAAGC | 1000 | 10 | 0.227 |
| t74 | Reverse | CTGAGACTTGCACATCGCAGCATCAACGACAAAAATAACAACTGATG | 1000 | 10 | 0.227 |
| t08 | Reverse | CTGAGACTTGCACATCGCAGCATATAAGAGATTCAAAGGGTCAGAC | 1000 | 10 | 0.227 |
| t82 | Reverse | CTGAGACTTGCACATCGCAGCGCTGGCGTATGTGTACACCT | 1000 | 10 | 0.227 |
| t85 | Reverse | CTGAGACTTGCACATCGCAGCAAGAATTTTAACACAAGGAGATCATC | 1000 | 10 | 0.227 |
| t96 | Reverse | CTGAGACTTGCACATCGCAGCGGGTGGTATCATGAGAATAGTTG | 1000 | 10 | 0.227 |
| t98 | Reverse | CTGAGACTTGCACATCGCAGCCAAACTTTTGGAGATTTCAAGATTATG | 1000 | 10 | 0.227 |
| t99 | Reverse | CTGAGACTTGCACATCGCAGCGGTCTTCCTAATCTTGAACCATC | 1000 | 10 | 0.227 |

***Supplementary Table 3.*** *Master mix composition for the primary multiplex ddPCR.*

| **Reagents** | **Final concentration** | **Volume** |
|---|---|---|
| NF ddH2O | | 6.5 µL |
| ddPCR Supermix for Probes (no dUTPs) | 1X | 11 µL |
| Primer Pool (35-plex) | 227nM | 0.5 µL |
| Template (DNA) | | 4 µL |
| Total | | 22 µL |

***Supplementary Table 4.*** *Cycling conditions of multiplex ddPCR.*

| **Step** | **Temperature** | **Time** | **Ramping** | **Cycles** |
|---|---|---|---|---|
| Initial Denaturation | 95°C | 10 min | - | 1 |
| Denaturation | 98°C | 30 sec | 2°C/s | 25 |
| Annealing/ | 58°C | 2 min | 2°C/s | |
| Extension | | | | |
| Enzyme deactivation | 98°C | 10 min | - | 1 |
| Hold | 4°C | ∞ | - | |

***Supplementary Table 5***. *PCR Primer sequences for AmpSeq library preparation (barcodes).*

| **Primer for Adapter PCR (XXXXXXXX=barcode)** | |
|---|---|
| Forward | AATGATACGGCGACCACCGAGATCTACACTCTTTCCCTACACGACGCTCTTCCGATCT**XXXXXXXX**GTGACCTATGAACTCAGGAGTC |
| Reverse | CAAGCAGAAGACGGCATACGAGATCGGTCTCGGCATTCCTGCTGAACCGCTCTTCCGATCT**XXXXXXXX**CTGAGACTTGCACATCGCAGC |

| **Forward barcode** | | **Reverse barcode** | |
|---|---|---|---|
| F01 | GCAACTGT | R01 | TCCGATCT |
| F02 | GAAGTACC | R02 | ACGAGAGA |
| F03 | GCCTTGAT | R03 | ATCTGTCC |
| F04 | CCAGGTTA | R04 | TGCTGCAA |
| F05 | GCTGACAA | R05 | CCACTAAG |
| F06 | TAGTGACG | R06 | TACCTTGC |
| F07 | GAGTTCGA | R07 | CGATCACA |
| F08 | AGAACCAC | R08 | ACCAAGTG |
| F09 | CTCCTCTA | R09 | CGGAAGAA |
| F10 | TTAACGCG | R10 | AATGGTGG |
| F11 | GGCATTCA | R11 | GGTTCCTT |
| F12 | CACTCTAG | R12 | GGACATTG |
| F13 | ATACAGGC | R13 | TCATACGG |
| F14 | AGTCGATC | R14 | CGTTATGC |
| F15 | TCTAGGAC | R15 | CTGGCATT |
| F16 | CACGAAGA | R16 | ACTCTGCA |
| F17 | AAGCACAG | R17 | GAACGGAA |
| F18 | AGCTTAGG | R18 | TTGGTCAC |
| F19 | CATAGCCA | R19 | GTGAACCT |
| F20 | TGGCAAGT | R20 | ATCGCGAA |
| F21 | GTCAGAAG | R21 | TGAGTGGA |
| F22 | GTTCCAGA | R22 | CTGATTGG |
| F23 | TCGCCTAA | R23 | GTTGTGTG |
| F24 | TGTGCAAG | R24 | GCGTAATC |

***Supplementary Table 6**. ENA accession numbers for WGS data from 198 monoclonal samples from MalariaGEN (Fws > 0.95) for relatedness inference and DAPC analysis.*

| SampleID | Origin | run_accessions |
|---|---|---|
| PR0118-C | Bangladesh | ERR216532 |
| PR0114-C | Bangladesh | ERR216644 |
| PR0159-C | Bangladesh | ERR404190 |
| PR0163-C | Bangladesh | ERR404203 |
| PR0132-C | Bangladesh | ERR404211 |
| PR0160-C | Bangladesh | ERR404230 |
| PR0141-C | Bangladesh | ERR404237 |
| PR0155-C | Bangladesh | ERR404241 |
| PR0131-C | Bangladesh | ERR404251 |
| PR0128-C | Bangladesh | ERR426105 |
| PH1492-C | Cambodia | ERR1172518 |
| PH1453-C | Cambodia | ERR1172522 |
| PH1611-C | Cambodia | ERR1172534 |
| PH1614-C | Cambodia | ERR1172535 |
| PH1499-C | Cambodia | ERR1172559 |
| PH1578-C | Cambodia | ERR1172579 |
| PH1634-C | Cambodia | ERR1172586 |
| PH1395-C | Cambodia | ERR905479 |
| PH1396-C | Cambodia | ERR905480 |
| QG0348-C | DRC | ERR1514555 |
| QG0357-C | DRC | ERR1514564 |
| QG0363-C | DRC | ERR1514570 |
| QG0380-C | DRC | ERR1514576 |
| QG0400-C | DRC | ERR1514583 |
| QG0430-C | DRC | ERR1514600 |
| QG0435-C | DRC | ERR1514602 |
| QG0307-C | DRC | ERR1514617 |
| QG0312-C | DRC | ERR1514619 |
| QG0325-C | DRC | ERR1514622 |
| QS0056-C | Ethiopia | ERR1035493 |
| QS0104-C | Ethiopia | ERR1035536 |
| QS0109-C | Ethiopia | ERR1045266 |
| QS0110-C | Ethiopia | ERR1045267 |
| QS0116-C | Ethiopia | ERR1045271 |
| QS0126-C | Ethiopia | ERR1045280 |
| QS0129-C | Ethiopia | ERR1045283 |
| QS0132-C | Ethiopia | ERR1045286 |
| QS0133-C | Ethiopia | ERR1045287 |
| QS0135-C | Ethiopia | ERR1045288 |
| QS0144-C | Ethiopia | ERR1045295 |
| QS0154-C | Ethiopia | ERR1106575 |
| QS0156-C | Ethiopia | ERR1106576 |
| QS0157-C | Ethiopia | ERR1106577 |
| QS0159-C | Ethiopia | ERR1106579 |
| QS0162-C | Ethiopia | ERR1106581 |
| QS0163-C | Ethiopia | ERR1106582 |
| QS0168-C | Ethiopia | ERR1106586 |
| QS0169-C | Ethiopia | ERR1106587 |
| QS0170-C | Ethiopia | ERR1106590 |
| QS0155-C | Ethiopia | ERR1106606 |
| PA0400-C | Gambia | ERR1106506 |
| PA0401-C | Gambia | ERR1106507 |
| PA0371-C | Gambia | ERR1106508 |
| PA0384-C | Gambia | ERR1106519 |
| PA0387-C | Gambia | ERR1106520 |

| | | |
|---|---|---|
| PA0396-C | Gambia | ERR1106528 |
| PA0398-C | Gambia | ERR1106530 |
| PA0417-C | Gambia | ERR1106543 |
| PA0618-C | Gambia | ERR1138790 |
| PA0655-C | Gambia | ERR1172503 |
| PF1069-C | Ghana | ERR636033 |
| PF1060-C | Ghana | ERR636054 |
| PF1037-C | Ghana | ERR636060 |
| PF1086-C | Ghana | ERR636068 |
| PF1027-C | Ghana | ERR636070 |
| PF1063-C | Ghana | ERR636072 |
| PF1028-C | Ghana | ERR636077 |
| PF1041-C | Ghana | ERR636086 |
| PF1102-C | Ghana | ERR636098 |
| PF1031-C | Ghana | ERR637303 |
| PA0151-C | Guinea | ERR055482 |
| PA0152-C | Guinea | ERR055493 |
| PA0208-C | Guinea | ERR059387 |
| PA0200-C | Guinea | ERR059396 |
| PA0207-C | Guinea | ERR059397 |
| PA0167-C | Guinea | ERR059407 |
| PA0189-C | Guinea | ERR059411 |
| PA0180-C | Guinea | ERR063529 |
| PA0144-C | Guinea | ERR063550 |
| PC0023-C | Kenya | ERR012317 |
| PC0003-02 | Kenya | ERR012359,ERR012498,ERR012257,ERR012334 |
| PC0068-C | Kenya | ERR012381,ERR012518 |
| PC0031-C | Kenya | ERR012425 |
| PC0064-C | Kenya | ERR012428 |
| PC0011-C | Kenya | ERR012475,ERR012330,ERR012446 |
| PC0047-C | Kenya | ERR012481 |
| PC0032-C | Kenya | ERR012494 |
| PC0063-C | Kenya | ERR012752,ERR012478 |
| PC0054-C | Kenya | ERR029083 |
| PC0057-C | Kenya | ERR029085 |
| PC0077-C | Kenya | ERR029088 |
| PC0046-C | Kenya | ERR029095 |
| PC0079-C | Kenya | ERR029104 |
| PC0075-C | Kenya | ERR029412 |
| PC0069-C | Kenya | ERR029413 |
| PC0053-C | Kenya | ERR126559 |
| PC0070-C | Kenya | ERR205934 |
| PC0104-C | Kenya | ERR205951 |
| PC0097-C | Kenya | ERR205958 |
| QE0400-C | Laos | ERR180058 |
| QE0454-C | Laos | ERR216572 |
| QE0455-C | Laos | ERR216573 |
| QE0457-C | Laos | ERR216574 |
| QE0475-C | Laos | ERR221487 |
| QE0419-C | Laos | ERR223057 |
| QE0430-C | Laos | ERR223075 |
| QE0438-C | Laos | ERR223081 |
| QE0440-C | Laos | ERR223083 |
| QE0405-Cx | Laos | ERR404175 |
| PT0031-C | Malawi | ERR054070 |
| PT0035-C | Malawi | ERR054071 |
| PT0041-C | Malawi | ERR054074 |
| PT0084-C | Malawi | ERR072008 |
| PT0079-C | Malawi | ERR108409 |
| PT0106-C | Malawi | ERR114390 |

| | | |
|---|---|---|
| PT0086-C | Malawi | ERR114394 |
| PT0149-C | Malawi | ERR216485 |
| PT0158-C | Malawi | ERR216580 |
| PT0170-C | Malawi | ERR216600 |
| PM0435-C | Mali | ERR662085 |
| PM0395-C | Mali | ERR662092 |
| PM0494-C | Mali | ERR666861 |
| PM0480-C | Mali | ERR666898 |
| PM0510-C | Mali | ERR666905 |
| PM0478-C | Mali | ERR666929 |
| PM0379-C | Mali | ERR670357 |
| PM0423-C | Mali | ERR670379 |
| PM0459-C | Mali | ERR670380 |
| PM0495-C | Mali | ERR670382 |
| QC0332-C | Myanmar | ERR1215361 |
| QC0342-C | Myanmar | ERR1215371 |
| QC0349-C | Myanmar | ERR1215378 |
| QC0351-C | Myanmar | ERR1215380 |
| QC0276-C | Myanmar | ERR1274866 |
| QC0277-C | Myanmar | ERR1274867 |
| QC0281-C | Myanmar | ERR1274870 |
| QC0285-C | Myanmar | ERR1274874 |
| QC0307-C | Myanmar | ERR1274889 |
| QC0328-C | Myanmar | ERR1274905 |
| QJ0007-C | Nigeria | ERR1172589 |
| QJ0008-C | Nigeria | ERR1172590 |
| QJ0018-C | Nigeria | ERR1172594 |
| QJ0160-C | Nigeria | ERR1172604 |
| QJ0164-C | Nigeria | ERR1172608 |
| QJ0167-C | Nigeria | ERR1172609 |
| QJ0169-C | Nigeria | ERR1172611 |
| QJ0172-C | Nigeria | ERR1172615 |
| QJ0174-C | Nigeria | ERR1172617 |
| QJ0175-C | Nigeria | ERR1214226 |
| PE0012-C | Tanzania | ERR018916 |
| PE0021-C | Tanzania | ERR022908 |
| PE0024-C | Tanzania | ERR022909 |
| PE0009-CW | Tanzania | ERR171645 |
| PE0105-C | Tanzania | ERR405252 |
| PE0134-C | Tanzania | ERR439523 |
| PE0124-C | Tanzania | ERR439528 |
| PE0125-C | Tanzania | ERR439531 |
| PE0137-C | Tanzania | ERR439532 |
| PE0130-C | Tanzania | ERR449903 |
| PE0305-C | Tanzania | ERR676469 |
| PE0273-C | Tanzania | ERR676472 |
| PE0294-C | Tanzania | ERR676482 |
| PE0314-C | Tanzania | ERR676485 |
| PE0207-C | Tanzania | ERR676491 |
| PE0255-C | Tanzania | ERR676497 |
| PE0393-C | Tanzania | ERR676499 |
| PE0336-C | Tanzania | ERR676508 |
| PE0492-C | Tanzania | ERR676514 |
| PE0493-C | Tanzania | ERR676534 |
| PE0547-C | Tanzania | ERR676542 |
| PE0213-C | Tanzania | ERR689161 |
| PE0437-C | Tanzania | ERR689171 |
| PE0479-C | Tanzania | ERR689173 |
| PE0426-C | Tanzania | ERR689180 |
| PE0173-C | Tanzania | ERR689186 |

| | | |
|---|---|---|
| PE0303-C | Tanzania | ERR689191 |
| PE0477-C | Tanzania | ERR689209 |
| PE0161-C | Tanzania | ERR689211 |
| PE0349-C | Tanzania | ERR696961 |
| PD0462-C | Thailand | ERR164726 |
| PD0521-C | Thailand | ERR216506 |
| PD0824-C | Thailand | ERR580376 |
| PD1161-C | Thailand | ERR596140 |
| PD1163-C | Thailand | ERR596151 |
| PD1204-C | Thailand | ERR689254 |
| PD1191-C | Thailand | ERR689264 |
| PD1189-C | Thailand | ERR689288 |
| PD1223-C | Thailand | ERR689308 |
| PD1220-C | Thailand | ERR689327 |
| PV0241-C | Vietnam | ERR126570 |
| PV0301-C | Vietnam | ERR164737 |
| PV0265-C | Vietnam | ERR171582 |
| PV0280-C | Vietnam | ERR180089 |
| PV0304-C | Vietnam | ERR216543 |
| PV0326-C | Vietnam | ERR221523 |
| PV0249-Cx | Vietnam | ERR388792 |
| PV0315-Cx | Vietnam | ERR404180 |
| PV0331-Cx | Vietnam | ERR404181 |

***Supplementary Table 7.*** *Estimates of multilocus linkage disequilibrium (mLD) for P. falciparum isolates from Gondar stratified by transmission seasons and Ziway. $I_A^S$ = standardized index of association. The Monte Carlo method (100,000 permutations) was used to test the significance of mLD.*

| **Population** | | **All infections** | | **Monoclonal infections** | |
|---|---|---|---|---|---|
| | | **n** | **$I_A^S$ (*p* value)** | **n** | **$I_A^S$ (*p* value)** |
| **Gondar** | | 63 | 0.0844 (<0.001) | 43 | 0.1061 (<0.001) |
| | **Wet season** | 31 | 0.0960 (<0.001) | 22 | 0.1202 (<0.001) |
| | **Dry season** | 32 | 0.0983 (<0.001) | 21 | 0.1381 (<0.001) |
| **Ziway** | | 17 | 0.1261 (<0.001) | 16 | 0.1371 (<0.001) |
| **All** | | 80 | 0.0725 (<0.001) | 59 | 0.0873 (<0.001) |

***Supplementary Table 8.*** *Frequency of mutations from drug resistance alleles in the population stratified by study site.*

| Gene | Mutation | Frequency% (n/N) | | *P* value [a] |
|---|---|---|---|---|
| | | **Gondar** | **Ziway** | |
| ***pfdhfr*** | N51I | 100% (152/152) | 100% (26/26) | NA |
| | **C59R** | **36.2% (55/152)** | **100% (26/26)** | **0.0005** |
| | S108N | 99.3% (151/152) | 100% (26/26) | 1 |
| | S108T | 0% (0/152) | 0% (0/26) | NA |
| | I164L | 0% (0/151) | 0% (0/26) | NA |
| ***pfdhps*** | K540E | 91.6% (141/154) | 100.0% (23/23) | 0.3553 |
| | A581G | 0% (0/154) | 0% (0/23) | NA |
| | A613T | 0% (0/154) | 0% (0/23) | NA |
| | A613S | 0% (0/154) | 0% (0/23) | NA |
| ***pfmdr1*** | N86Y | 3.3% (5/152) | 0% (0/26) | 0.6122 |
| | Y184F | 98.7% (156/158) | 100.0% (26/26) | 1 |
| ***pfmdr2*** | **I492V** | **39.7% (60/151)** | **12.0% (3/25)** | **0.0125** |
| | T484I | 0% (0/151) | 0% (0/25) | NA |
| ***pfk13*** | I543T | 0% (0/151) | 0% (0/26) | NA |
| | R539T | 0% (0/151) | 0% (0/26) | NA |
| | G538V | 0% (0/151) | 0% (0/26) | NA |
| | N537I | 0% (0/151) | 0% (0/26) | NA |
| | P527H | 0% (0/151) | 0% (0/26) | NA |
| | Y493H | 0% (0/151) | 0% (0/26) | NA |
| | A481V | 0% (0/52) | 0% (0/26) | NA |

[a] All *P* values were 2-tailed and computed using the Pearson's Chi-squared test with simulated *P* value (based on 2000 replicates) for comparing the proportion of mutant alleles.

***Supplementary Table 9.*** *Mean pairwise IBD and proportion of related infection pairs between samples from both study sites (Gondar and Ziway) and several African countries. P values of each comparison were calculated by permutation test (100,000 permutations).*

| **Pairs (with Gondar & Ziway)** | **n pairs** | **Mean IBD** | ***P value*** | **Proportion related infections** | ***P value*** |
|---|---|---|---|---|---|
| **Southeast Asia** | | | | | |
| **Bangladesh** | 1,870 | 0.0010 | >0.05 | 0.00% | >0.05 |
| **Myanmar** | 1,870 | 0.0021 | >0.05 | 0.11% | >0.05 |
| **Thailand** | 1,870 | 0.0029 | >0.05 | 0.05% | >0.05 |
| **Laos** | 1,870 | 0.0008 | >0.05 | 0.00% | >0.05 |
| **Cambodia** | 1,683 | 0.0010 | >0.05 | 0.00% | >0.05 |
| **Vietnam** | 1,683 | 0.0012 | >0.05 | 0.00% | >0.05 |
| **West Africa** | | | | | |
| **Gambia** | 1,870 | 0.0037 | >0.05 | 0.16% | >0.05 |
| **Guinea** | 1,683 | 0.0057 | >0.05 | 0.71% | >0.05 |
| **Mali** | 1,870 | 0.0037 | >0.05 | 0.21% | >0.05 |
| **Ghana** | 1,870 | 0.0044 | >0.05 | 0.59% | >0.05 |
| **Nigeria** | 1,870 | 0.0058 | >0.05 | 0.37% | >0.05 |
| **Central Africa** | | | | | |
| **DRC** | 1,870 | 0.0096 | >0.05 | 0.80% | >0.05 |
| **East Africa** | | | | | |
| **Malawi** | 1,870 | 0.0061 | >0.05 | 0.21% | >0.05 |
| **Tanzania** | 5,610 | 0.0088 | >0.05 | 0.77% | >0.05 |
| **Kenya** | | 0.0065 | >0.05 | 0.40% | >0.05 |
| **Horn of Africa** | | | | | |
| **Ethiopia (MalariaGEN)** | 3,927 | 0.0893 | **0.0009** | 11.71% | 0.1010 |

***Supplementary Table 10**. Mean pairwise IBD and proportion of related infection pairs between samples from Gondar and several African countries. P values of each comparison were calculated by permutation test (100,000 permutations).*

| Pairs (with Gondar) | n pairs | Mean IBD | *P value* | Proportion related infections | *P value* |
|---|---|---|---|---|---|
| **Southeast Asia** | | | | | |
| **Bangladesh** | 1,590 | 0.0009 | >0.05 | 0.00% | >0.05 |
| **Myanmar** | 1,590 | 0.0011 | >0.05 | 0.06% | >0.05 |
| **Thailand** | 1,590 | 0.0008 | >0.05 | 0.06% | >0.05 |
| **Laos** | 1,590 | 0.0004 | >0.05 | 0.00% | >0.05 |
| **Cambodia** | 1,431 | 0.0007 | >0.05 | 0.00% | >0.05 |
| **Vietnam** | 1,431 | 0.0006 | >0.05 | 0.00% | >0.05 |
| **West Africa** | | | | | |
| **Gambia** | 1,590 | 0.0029 | >0.05 | 0.19% | >0.05 |
| **Guinea** | 1,431 | 0.0045 | >0.05 | 0.49% | >0.05 |
| **Mali** | 1,590 | 0.0032 | >0.05 | 0.25% | >0.05 |
| **Ghana** | 1,590 | 0.0046 | >0.05 | 0.69% | >0.05 |
| **Nigeria** | 1,590 | 0.0058 | >0.05 | 0.44% | >0.05 |
| **Central Africa** | | | | | |
| **DRC** | 1,590 | 0.0098 | >0.05 | 0.94% | >0.05 |
| **East Africa** | | | | | |
| **Malawi** | 1,590 | 0.0062 | >0.05 | 0.25% | >0.05 |
| **Tanzania** | 4,770 | 0.0086 | >0.05 | 0.75% | >0.05 |
| **Kenya** | 3,180 | 0.0064 | >0.05 | 0.41% | >0.05 |
| **Horn of Africa** | | | | | |
| **Ethiopia (MalariaGEN)** | 3,339 | 0.0853 | **0.0032** | 10.84% | **0.0036** |
| **Ziway** | 4,452 | 0.1082 | **<0.0001** | 14.53% | **0.0002** |

***Supplementary Table 11.*** *Mean pairwise IBD and proportion of related infection pairs between samples from Ziway and several African countries. P values of each comparison were calculated by permutation test (100,000 permutations).*

| Pairs (with Ziway) | n pairs | Mean IBD | *P value* | Proportion related infections | *P value* |
|---|---|---|---|---|---|
| **Southeast Asia** | | | | | |
| **Bangladesh** | 280 | 0.0013 | >0.05 | 0.00% | >0.05 |
| **Myanmar** | 280 | 0.0075 | >0.05 | 0.36% | >0.05 |
| **Thailand** | 280 | 0.0144 | >0.05 | 0.00% | >0.05 |
| **Laos** | 280 | 0.0032 | >0.05 | 0.00% | >0.05 |
| **Cambodia** | 252 | 0.0025 | >0.05 | 0.00% | >0.05 |
| **Vietnam** | 252 | 0.0042 | >0.05 | 0.00% | >0.05 |
| **West Africa** | | | | | |
| **Gambia** | 280 | 0.0078 | >0.05 | 0.00% | >0.05 |
| **Guinea** | 252 | 0.0121 | >0.05 | 1.98% | >0.05 |
| **Mali** | 280 | 0.0063 | >0.05 | 0.00% | >0.05 |
| **Ghana** | 280 | 0.0037 | >0.05 | 0.00% | >0.05 |
| **Nigeria** | 280 | 0.0055 | >0.05 | 0.00% | >0.05 |
| **Central Africa** | | | | | |
| **DRC** | 280 | 0.0085 | >0.05 | 0.00% | >0.05 |
| **East Africa** | | | | | |
| **Malawi** | 280 | 0.0053 | >0.05 | 0.00% | >0.05 |
| **Tanzania** | 840 | 0.0098 | >0.05 | 0.83% | >0.05 |
| **Kenya** | 560 | 0.0068 | >0.05 | 0.36% | >0.05 |
| **Horn of Africa** | | | | | |
| **Ethiopia (MalariaGEN)** | 588 | 0.1120 | **0.0008** | 16.67% | **0.00362** |
| **Gondar** | 4,452 | 0.1082 | **<0.0001** | 14.53% | **0.0002** |

### SUPPLEMENTARY FIGURES

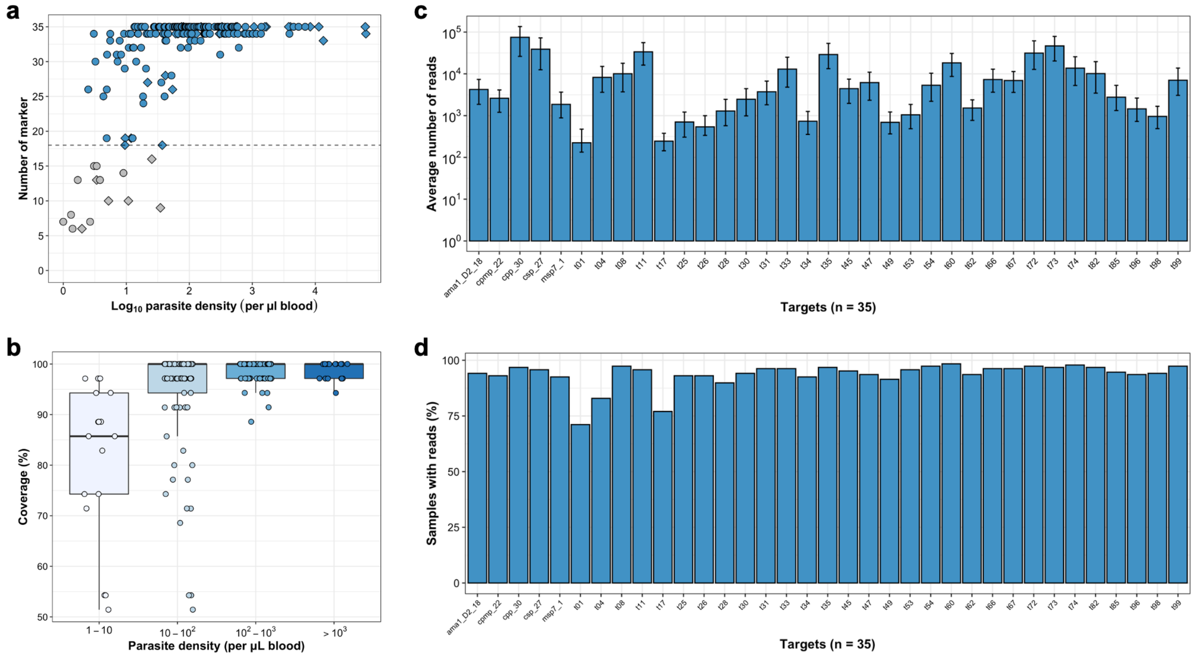


***Supplementary Figure 1.*** *Evenness and coverage of multiplexed amplicon sequencing of microhaplotypes (n=28) and drug resistance loci (n=7).* ***A****, Coverage of microhaplotype loci and drug resistance targets by parasite density in 168 DBS samples from Gondar. 159 samples with data in ≥18 loci (>50% coverage; dashed line, in blue) were included for further analyses.* ***B****, Boxplot summarizing the coverage of microhaplotype loci and drug resistance targets by parasite density in 159 DBS samples with (>50% coverage.* ***C****, Average number of reads per target per sample. The median (bars) and interquartile range (error bars) are shown. Note that the y-axis is on a log10-scale.* ***D****, Number of samples (%) of the 187 samples included with ≥50 reads per marker.*

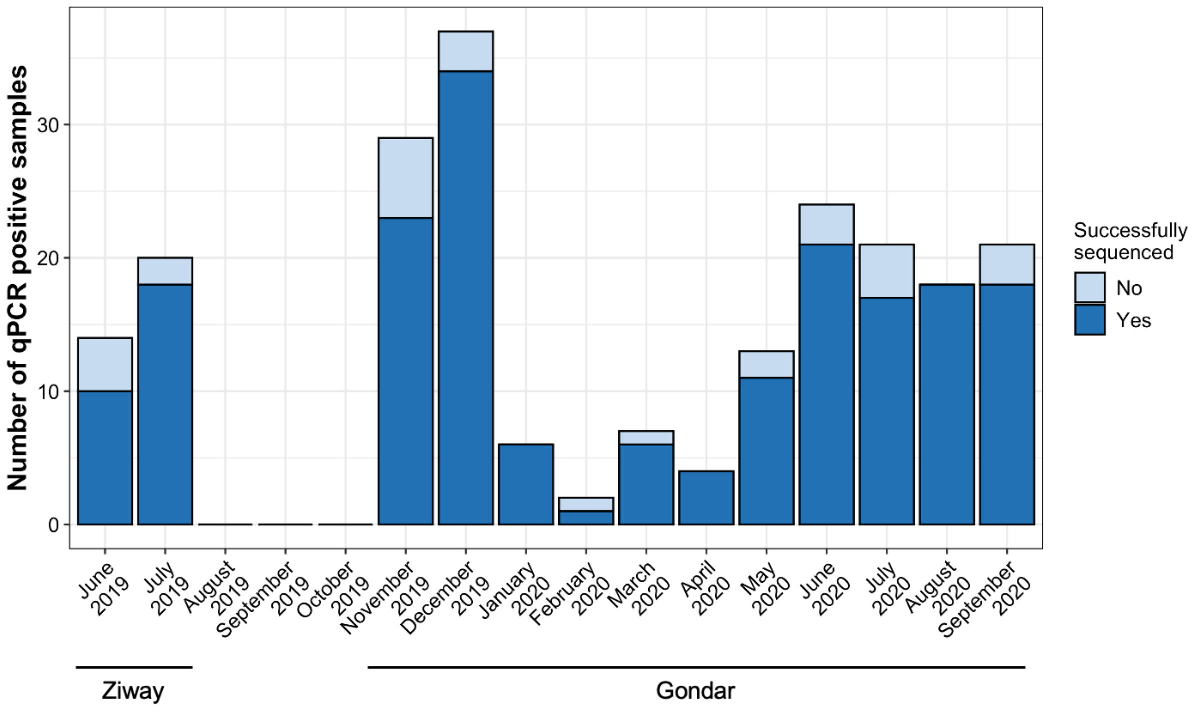


***Supplementary Figure 2.*** *Number of P. falciparum positive samples by qPCR (n=202) with ≥1 parasite/μL across different months. Colors indicate the number of successfully sequenced samples (n=187; >50% coverage) for each month. Sample collection in Gondar did not occur on all days in January and February. Collection periods for both studies are indicated.*

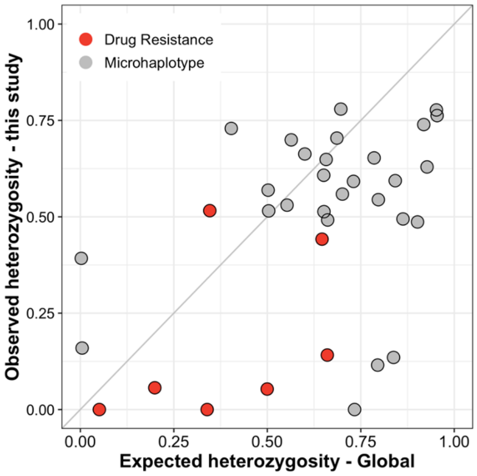


***Supplementary Figure 3.*** *Comparison of expected heterozygosity of microhaplotypes and drug resistance loci in the global population and the observed heterozygosity in this study (Gondar and Ziway samples).*

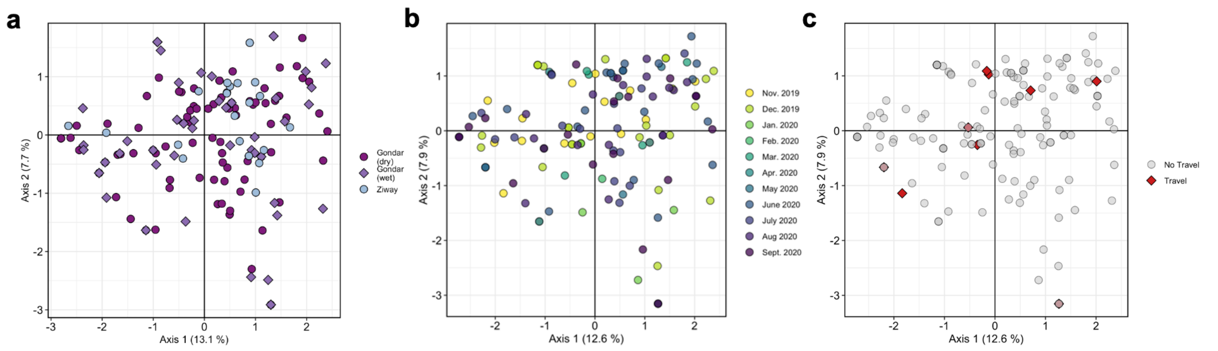


***Supplementary Figure 4.*** *Principal components analysis (PCA) using allele frequency of the 28 microhaplotypes from dominant alleles.* ***A****, Samples from Gondar and Ziway. The first two PCA components are shown (explaining 13.1% and 7.7% of variation in the dataset, respectively).* ***B,C*** *Samples from Gondar only, colored (****B****) months and (****C****) travel history. The first two PCA components are shown (explaining 12.6% and 7.9% of variation in the dataset, respectively).*

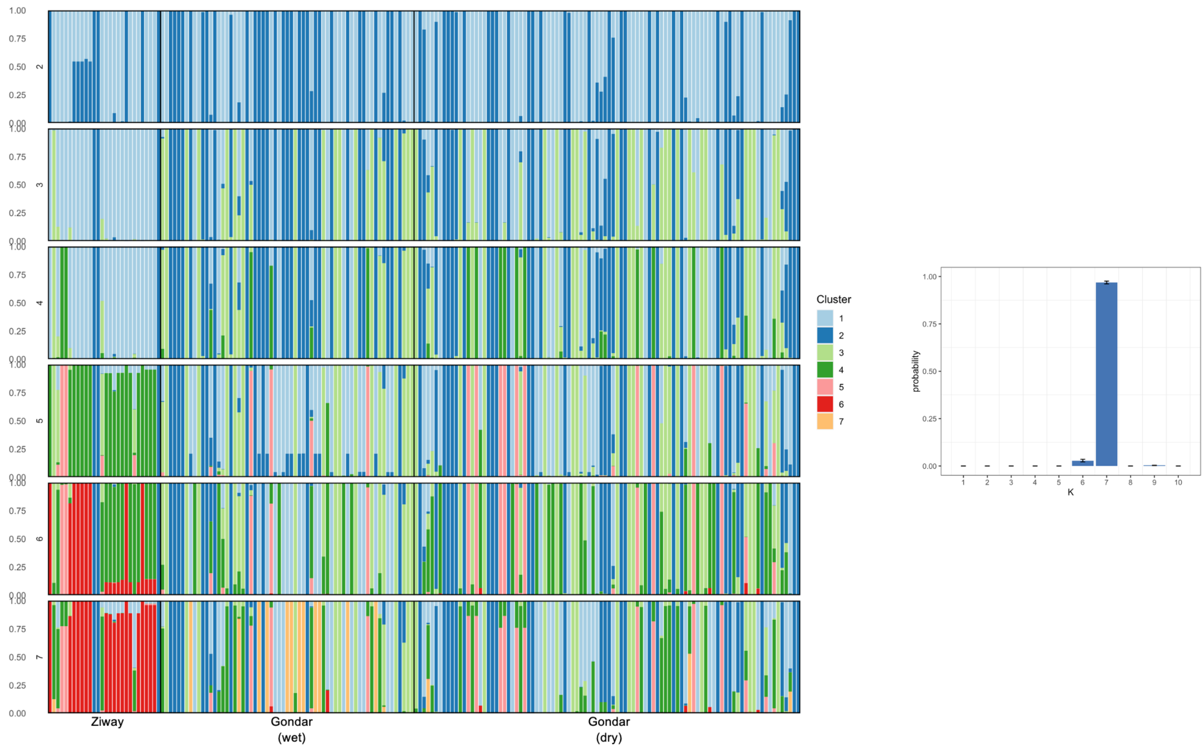


***Supplementary Figure 5****. Population cluster analysis of P. falciparum microhaplotypes from dominant alleles in Gondar and Ziway. Individual ancestry coefficients and different K values (K=2-7) are shown as inferred by rmaverick. Each vertical bar represents an individual haplotype and its membership to the eight cluster are defined by the different colors. Black borders separate the study sites. Samples are ordered chronologically. Barplot showing optimum K value (K=7).*

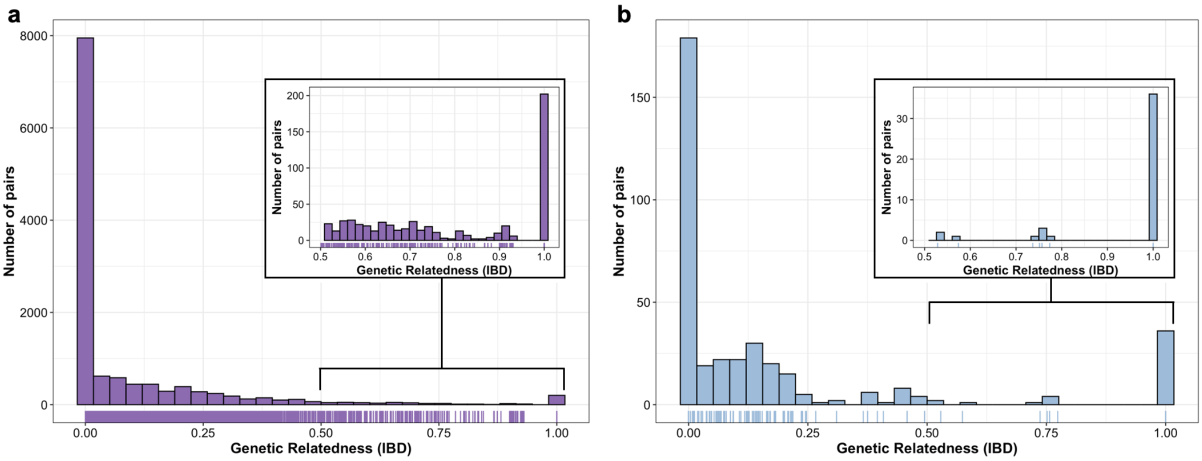


***Supplementary Figure 6.*** *Histogram of pairwise IBD. Pairwise IBD between (**a**) Gondar samples (n=12,561) and (**b**) Ziway samples (n=378), estimated by dcifer. Inset shows the heavy tail of the distribution, with some pairs of samples having IBD = 1.*

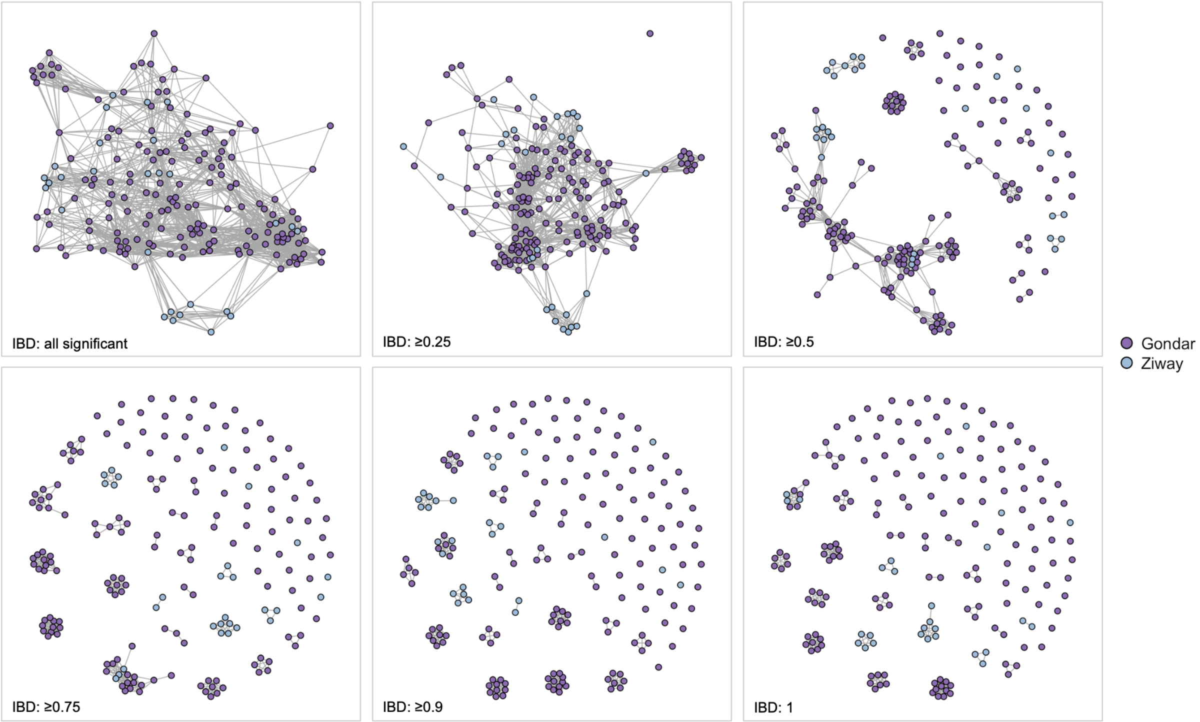


***Supplementary Figure 7.*** *Network analysis of pairwise relatedness visualized using different thresholds of IBD and colored by study site. Nodes correspond to indicated IBD threshold.*

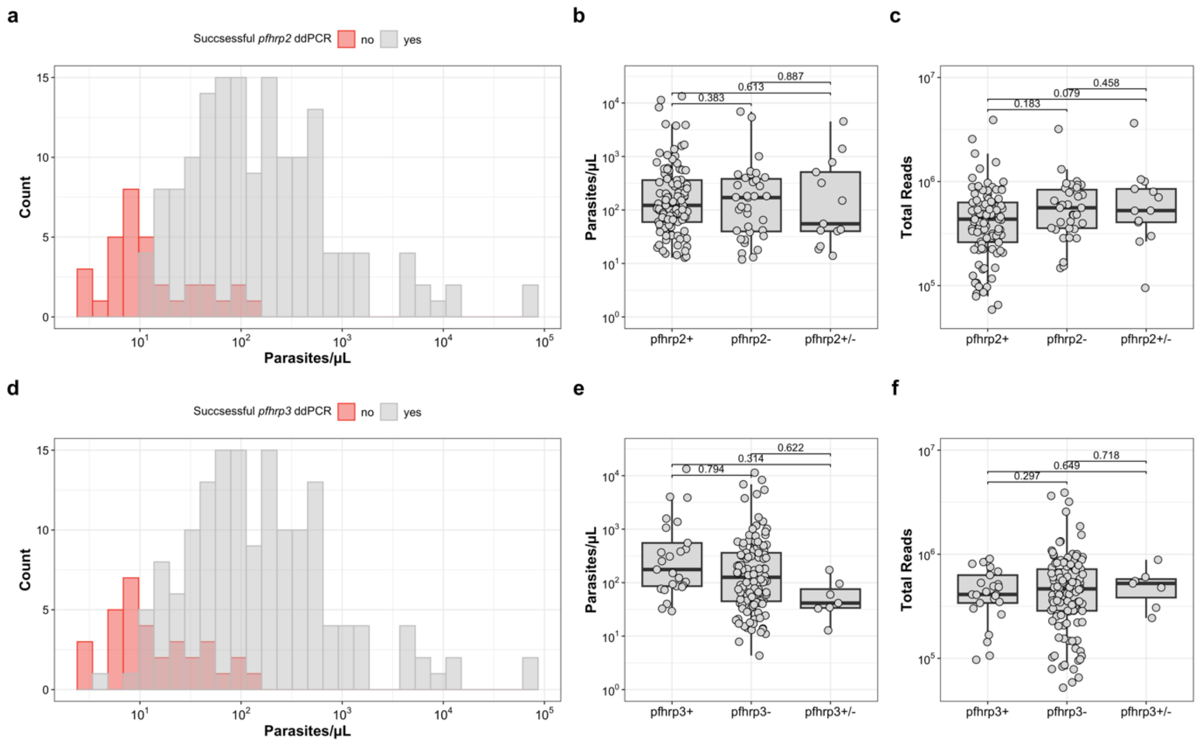


***Supplementary Figure 8****. Successful ddPCR pfhrp2 or pfhrp3 deletion status calls versus qPCR parasite density and the number of sequencing reads. Comparison of pfhrp2 (**A**) or pfhrp3 (**D**) ddPCR results versus varATS qPCR parasite densities. Samples were considered as successful if they have at least 10 tRNA copies in the ddPCR. Parasite densities of successfully typed samples by qPCR by pfhrp2 (**B**) or pfhrp3 (**E**) deletion status. Total number of reads of successfully typed samples by pfhrp2 (**C**) or pfhrp3 (**F**) deletion status. Differences among parasite densities or number of reads were assessed by Student t test.*

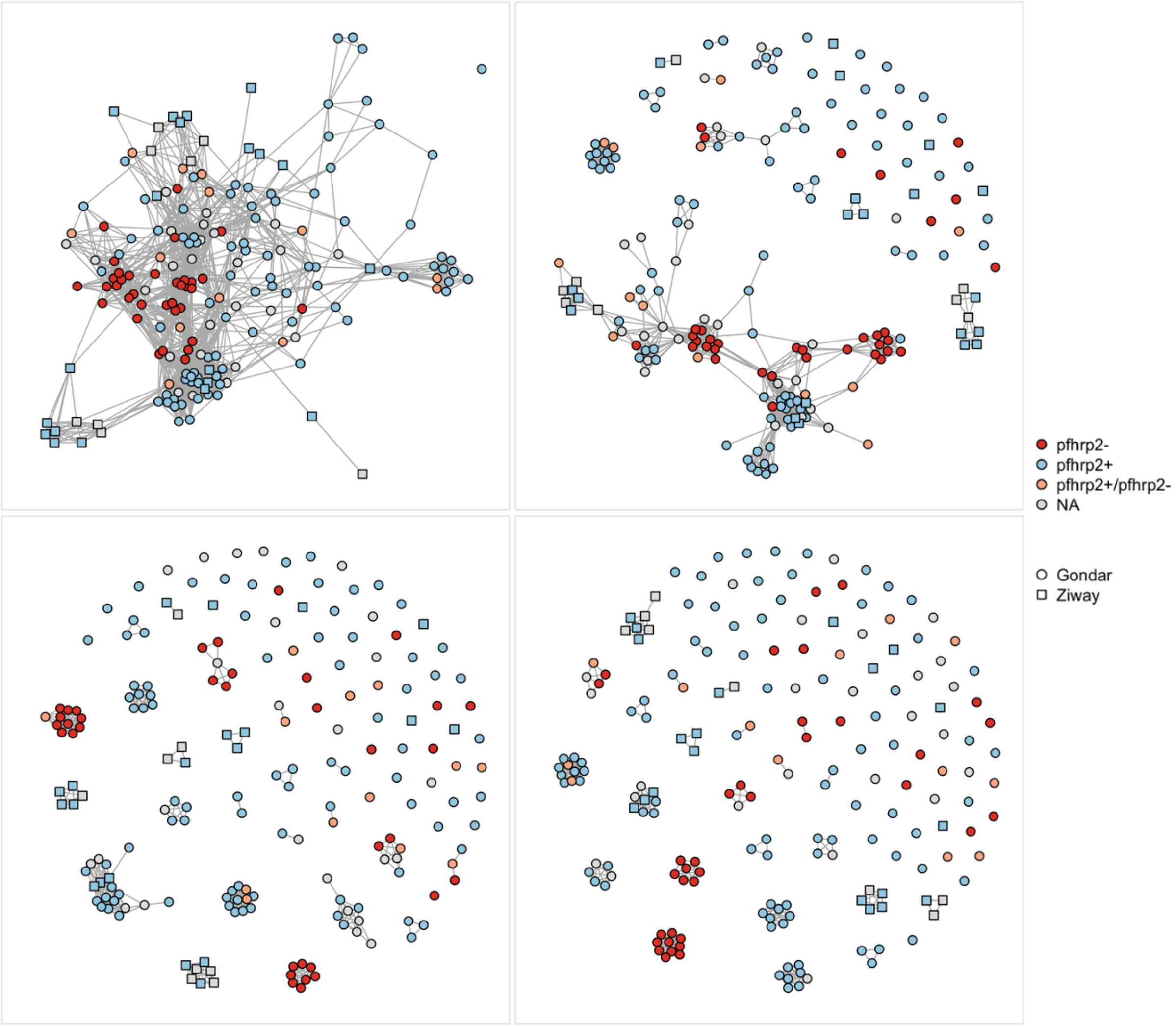


***Supplementary Figure 9.*** *Network analysis of pairwise relatedness visualized using different thresholds of IBD and colored by pfhrp2 deletion status. Nodes correspond to indicated IBD threshold. Circles (Gondar) and squared (Ziway) indicate study sites.*

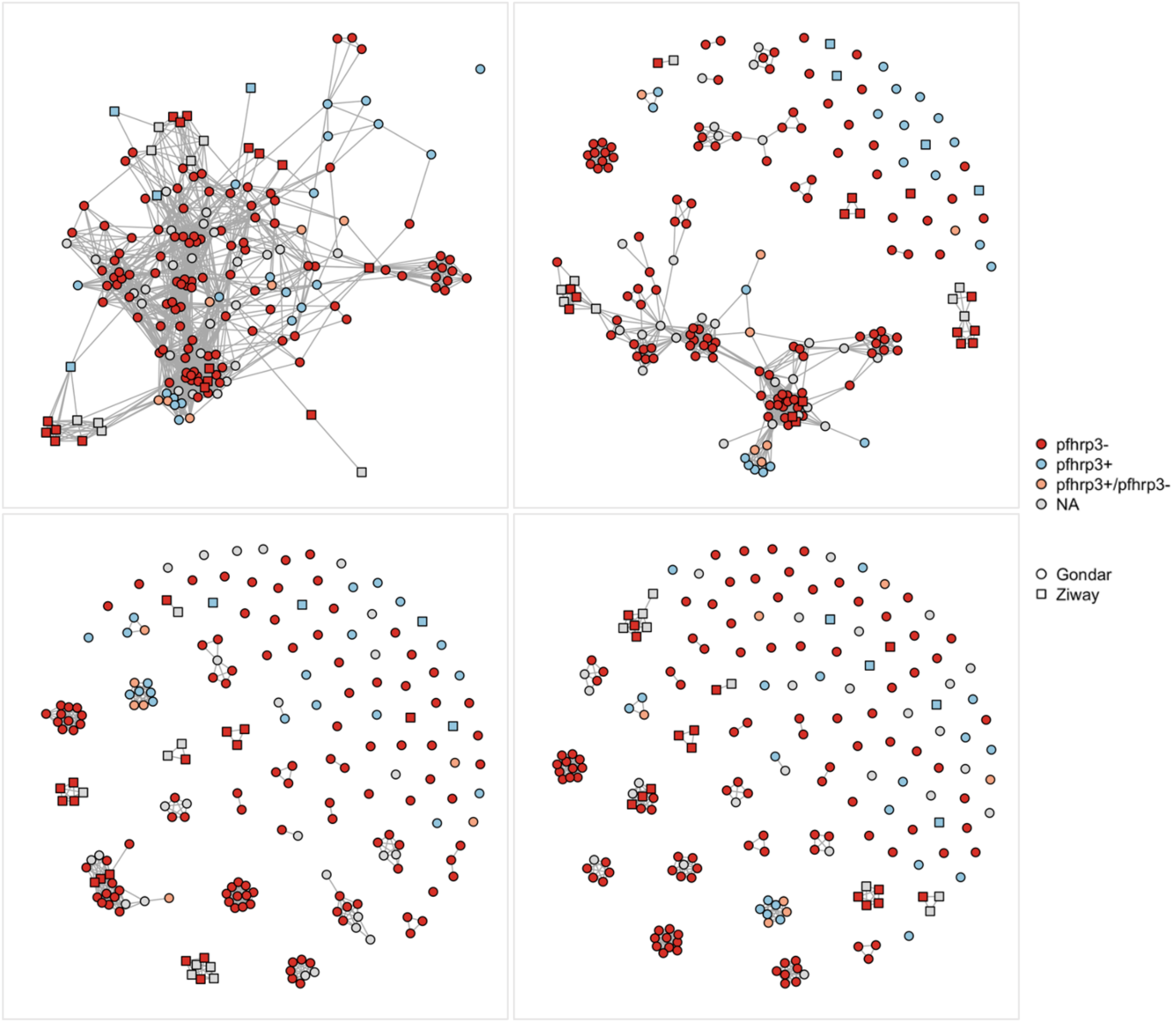


***Supplementary Figure 10.*** *Network analysis of pairwise relatedness visualized using different thresholds of IBD and colored by pfhrp3 deletion status. Nodes correspond to indicated IBD threshold. Circles (Gondar) and squared (Ziway) indicate study sites.*

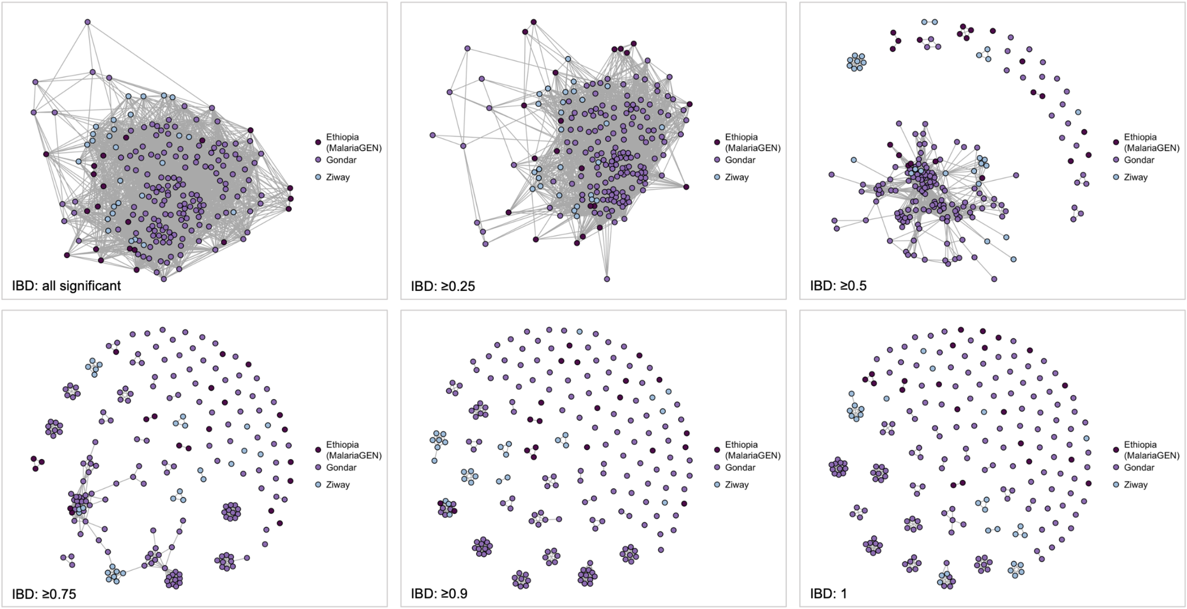


***Supplementary Figure 11.*** *Network analysis of pairwise relatedness visualized using different thresholds of IBD and colored by sample origin. Nodes correspond to indicated IBD threshold.*

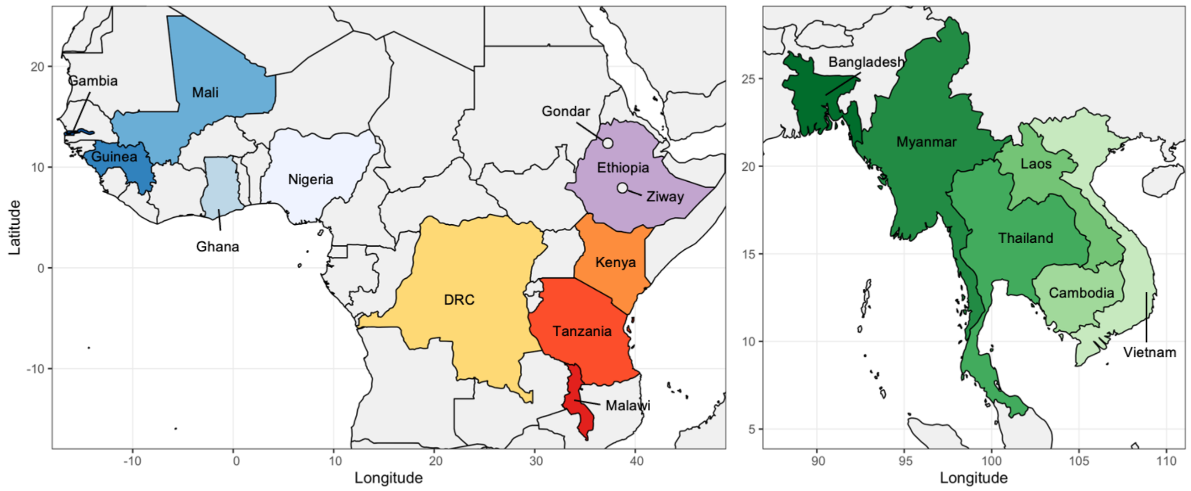


***Supplementary Figure 12***. *Maps showing countries with WGS samples from the MalariaGEN database that were included in our analysis. Left: Map of Africa. 140 samples were included from 10 different countries. The locations of our study sites are indicated by the white circles. Right: Map of Southeast Asia. 58 samples were included from 6 different countries.*
